# Mobile genetic elements shape the evolution of wild bacterial pangenomes in the face of biotic and abiotic selective pressures in the soil

**DOI:** 10.1101/2025.10.22.684009

**Authors:** Angeliqua P. Montoya, Hanna Kehlet-Delgado, Kyson T. Jensen, Joel S. Griffitts, Stephanie S. Porter

**Affiliations:** School of Biological Sciences, Washington State University, Vancouver, Washington, 98686; Department of Plant Pathology, Washington State University, Pullman, WA 99164; Department of Microbiology and Molecular Biology, Brigham Young University, Provo, Utah, 84602

## Abstract

Mobile genetic elements (MGEs) provide a critical reservoir of diversity that potentiates bacterial local adaptation. However, the degree to which natural populations of MGEs and the chromosomes that host them share congruent or divergent evolutionary histories remains an open question. MGE and host chromosome phylogenies could show congruence or independence, depending on rates of MGE gain and loss, costs and benefit of MGE carriage, compatibility, and gene flow. We investigated phylogenetic congruence for two critical niche-expanding MGEs and host chromosomes using 302 genomes of wild symbiotic nitrogen-fixing *Mesorhizobium* bacteria. The MGEs comprise: 1) a nickel resistance island (NRI), which confers adaptation to nickel-enriched serpentine soils across replicated natural populations, and 2) a more extensive symbiosis island (SI), which enables symbiotic nitrogen fixation with a host legume. We demonstrate that the NRI, like the SI, is transmissible and its acquisition confers nickel adaptation. NRI acquisition often incurs no detectable cost, consistent with conditional neutrality during MGE-mediated adaptation. Phylogenies for the NRI, SI, and chromosome are congruent across wild strains, consistent with a history of vertical co-inheritance and/or MGE transmission primarily among close relatives. This congruence does not appear to result from barriers to gene flow, as diverse MGE and host chromosome haplotypes are broadly distributed and co-occur. Thus, despite ample ecological opportunity for admixture, distant relatives rarely transmitted MGEs, consistent with selection to maintain co-adapted genomic compartments. The SI has stronger phylogenetic congruence with the chromosome than does the smaller NRI, supporting the hypothesis that MGE complexity limits horizontal transfer among more divergent lineages. However, rare horizontal NRI transmission events among distant relatives and NRI loss appear to occur when haplotypes migrate across environments. Thus, environmentally dependent selection on migrants may periodically disrupt co-adapted MGE and chromosome haplotypes and serve as an engine of diversity.

## INTRODUCTION

The bacterial mobilome, which consists of all the mobile elements in a bacterial genome, plays a key role in bacterial fitness, adaptation, and niche expansion. Mobile genetic elements (MGEs), comprise discrete sequences that can be transmitted vertically during cell division or horizontally among lineages. MGEs include integrative and conjugative elements (ICEs), plasmids, and prophages (1,2). MGEs can confer novel gene functions and potentiate gene gain and loss, which can enable adaptation to new environments and the emergence of divergent populations, profoundly altering bacterial fitness, evolutionary dynamics, and diversity (2–6). How MGE exchange shapes bacterial adaptation and diversity across natural landscapes remains an open evolutionary question (1,7).

Both environmental and intragenomic selection shape patterns of MGE transmission (8). Across variable environments, MGE acquisition may be favored by local selection, with MGE loss favored in the absence of such selection. Concurrently, natural selection may favor specialization via acquisition of MGEs that are highly compatible with the recipient genome such that they bear co-adapted gene complexes that incur few negative epistatic effects (9–11). On the other hand, generalist MGEs may evolve to function well across diverse recipient genomes (4,12,13). Gene connectivity (i.e., the number of predicted protein-protein interactions in which a gene participates) is predicted to limit HGT (the “complexity hypothesis” sensu (14,15)). Generalist MGEs could arise due to co-transmission of essential genes for MGE function and the evolution of low gene connectivity between MGEs and recipient genomes (16). Furthermore, MGE transmission requires that donor/recipient lineages co-occur, and so the geographical distribution of particular haplotypes may be indicative of constraints to transmission and thus haplotype stability. Understanding how tensions between environmental pressures, transmission opportunities, and haplotype stability shape evolutionary trajectories for genomic compartments is a critical next step for understanding bacterial evolution in natural landscapes (1).

MGEs and recipient chromosomes can each be evaluated phylogenetically. When compared, these phylogenies may range from showing complete congruence to complete evolutionary independence. MGE-chromosome phylogenetic congruence may arise if compensatory mutations ameliorate previously incompatible genetic interactions. This process of MGE-chromosome co-adaptation may result in selective sweeps for high-fitness MGE/chromosome combinations (3,11,17). This dynamic could promote vertical MGE transmission or horizontal transmission among very close relatives (17–19) and result in a shared evolutionary history among genomic compartments (20,21). Furthermore, if locally adaptive MGEs incur no cost under non-local environments, this can promote stable vertical transmission and infrequent MGE loss (8,22). At the extreme, an MGE could be ‘domesticated’ and assimilated into the vertically transmitting chromosome with concomitant loss of horizontal transfer capability. However, domesticated haplotypes may suffer from more limited evolvability in the face of new environmental pressures (23,24).

In contrast, if adaptive MGEs have high horizontal transmission rates and/or incur little genetic incompatibility among diverse hosts, they may be relatively transient in a host (3,18). For example, antibiotic resistance genes are transmitted and acquired by diverse bacteria under selective pressure but purged in antibiotic-free environments (2,13,18). Here, MGEs and recipient genome compartments would evolve independently and show dissimilar phylogenies. Ultimately, cophylogenetic patterns at intermediate points between these possible signatures of high vs low rates of horizontal transfer may emerge as an evolutionary compromise, allowing some coevolution among genomic compartments but also influxes of genetic variation (25,26).

These potential scenarios are rarely examined quantitatively in wild bacterial populations across natural landscapes, especially for MGEs critical to adaptation. We can think of MGEs and potential host cells as interacting taxa. Phylogenetic congruence—the topological matching of phylogenies—measures codependence between interacting taxa which can arise from co-transmission or preferential transmission among close relatives (27). Cophylogenetic signal—the tendency for closely related members of one taxon to interact with closely related members of another taxon—will always present under phylogenetic congruence but can also arise when interacting taxa evolve independently and lack phylogenetic congruence (27). Investigating phylogenetic congruence and cophylogenetic signal among genomic compartments, especially adaptive MGEs across variable environments, is critical to evaluate evolutionary constraints on bacterial adaptation in nature.

We compare the evolutionary history of adaptive MGEs and recipient chromosomes in rhizobia bacteria, which reside in soil but may become nitrogen-fixing symbionts on leguminous host plants (28). In *Mesorhizobium*, the genes required for symbiosis and nitrogen fixation with a host legume reside on a ∼500 kb symbiosis island (SI). The SI is an ICE that bears genes required for conjugation and integration into a conserved Phe-tRNA gene in the bacterial chromosome (29–31). The SI is transmissible via conjugation in the wild and *in vitro*, and SI acquisition can confer functional symbiosis in some cases (32–34). Genes in the SI, such as *nif* genes that underlie nitrogen fixation, interact with recipient genomes and with host plant genes during symbiosis (35–37), consistent with characterization of the SI as a complex MGE with high gene connectivity within the genome. Genes in the SI, such as *nod* genes that underlie molecular interactions necessary for nodule initiation, determine compatibility with host plant species and thus host species impose differential selection on genetic variants in the SI (38–40).

*Mesorhizobium* can also harbor a nickel resistance island (NRI) bearing the *nre* operon, which confers nickel tolerance, and thus adaptation to serpentine soil, which is naturally enriched in toxic heavy metals, particularly nickel (Ni) (41–43). The NRI is ∼54 kb in reference strain C089B, where it bears molecular features consistent with an integrated and mobilizable element (IME), such as insertion at a tRNA gene, and a predicted requirement of conjugative functions in *trans* from another system, though NRI transmission has not previously been documented experimentally (42). In a pattern that has evolved repeatedly at sites across the landscape, NRI occurs at high frequency and with *nre* alleles that confer high Ni resistance in strains residing in Ni-enriched serpentine soils; but is it is usually absent or confers low Ni resistance in strains residing in non-serpentine soils, consistent with soil chemistry imposing selection on NRI presence/absence variation and allelic diversity (42). Within the NRI, only a few genes within the *nre* operon are necessary and sufficient to confer nickel tolerance, suggesting that NRI loci have low gene connectivity within the genome. Expression of this operon is inducible by Ni. It contains *nreX* and *nreY* which encode putative Ni efflux pumps, and *nreA*, a CsoR/RcnR-family transcriptional repressor that is responsive to exogenous Ni (44).

We investigate the contemporary genomics and phylogenomics of adaptive MGEs that expand the rhizobia niche across abiotic and biotic axes. We focus on NRI and SI elements in *Mesorhizobium* strains extracted from root nodules of *Acmispon* legumes. To investigate how transmission dynamics for these adaptive MGEs shapes wild bacterial evolution over short and long timescales, we ask: 1) Is the NRI transmissible among *Mesorhizobium* strains, and does transfer confer resistance to nickel? We reveal NRI mobility and functional transfer in laboratory studies and compare this to historic patterns of MGE transmission across natural landscapes by asking, 2) To what degree is the core NRI phylogeny congruent with that of the chromosome and the SI? We find that the NRI, a locus for abiotic adaptation, shows weaker phylogenetic congruence with the chromosome than does the SI, a locus that underlies biotic adaptation. To investigate further we ask, 3) Does MGE distribution reflect environmentally variable selection? We find patterns of phylotype functional differentiation and ancestral trait state evolution consistent with MGE evolution in response to variable selection. We thus integrate laboratory-based genetic manipulations with phylogenomic analyses to shed light on how MGEs shape wild bacterial pangenomes in the face of biotic and abiotic selective pressures across the landscape.

## RESULTS

We examined a wild *Mesorhizobium* population associated with native grassland legumes in Western USA described in Kehlet Delgado et al. (2024). Briefly, *Mesorhizobium* bacteria were isolated from *Acmispon wrangelianus* and *A. brachycarpus* root nodules collected along a 12 m transect at each of 56 sites. Sites spanned 15 natural reserves across 1000 km in latitude from Southern California to Southern Oregon, USA. Sites encompassed physiologically stressful serpentine soils enriched in toxic nickel as well as more fertile, non-serpentine soils, and soil nickel enrichment was measured at each site. In total, 302 genomes (286 draft genomes via Illumina and 16 complete genomes via PacBio) were characterized (42). For each strain, ecological factors were recorded at the time of isolation, such as the host plant species and soil type (serpentine or non-serpentine soil), and nickel tolerance was measured *in vitro* as the minimum inhibitory concentration of Ni (MIC) (42).

### NRI is transmissible as an IME

To test horizontal transmission of the NRI, we tracked transfer from an NRI+ donor to recipient strains by targeted insertion of selectable markers into the NRI, followed by genomic sequencing. We refer to bacterial haplotypes as donors (capable of transferring an MGE), recipients (capable of acquiring an MGE), or transconjugants (accepted a novel MGE). The NRI of reference donor strain C089B (Table S1) was transformed with a synthetic plasmid bearing markers for neomycin (Nm) resistance and beta-glucuronidase (*GUS*), which cleaves a substrate to stain cells blue in the presence of X-gluc (5-Bromo-4-chloro-3-indoxyl beta-D-glucuronide). The plasmid contains a region of homology with the opine dehydrogenase gene, *ODH*, such that it can recombine into adjacent intergenic space in the NRI without disrupting gene function (see Supplemental Information for detail; Figure 1a; Figure S1). The C089B NRI is inserted at a Met-tRNA gene and ends in a 14-bp direct repeat of the end of the tRNA gene (42). Six *Mesorhizobium* recipients were mated with the donor, including four strains that initially lacked the NRI and two strains that already possessed copies of the NRI (Table S1; see Methods).

**Figure 1.**
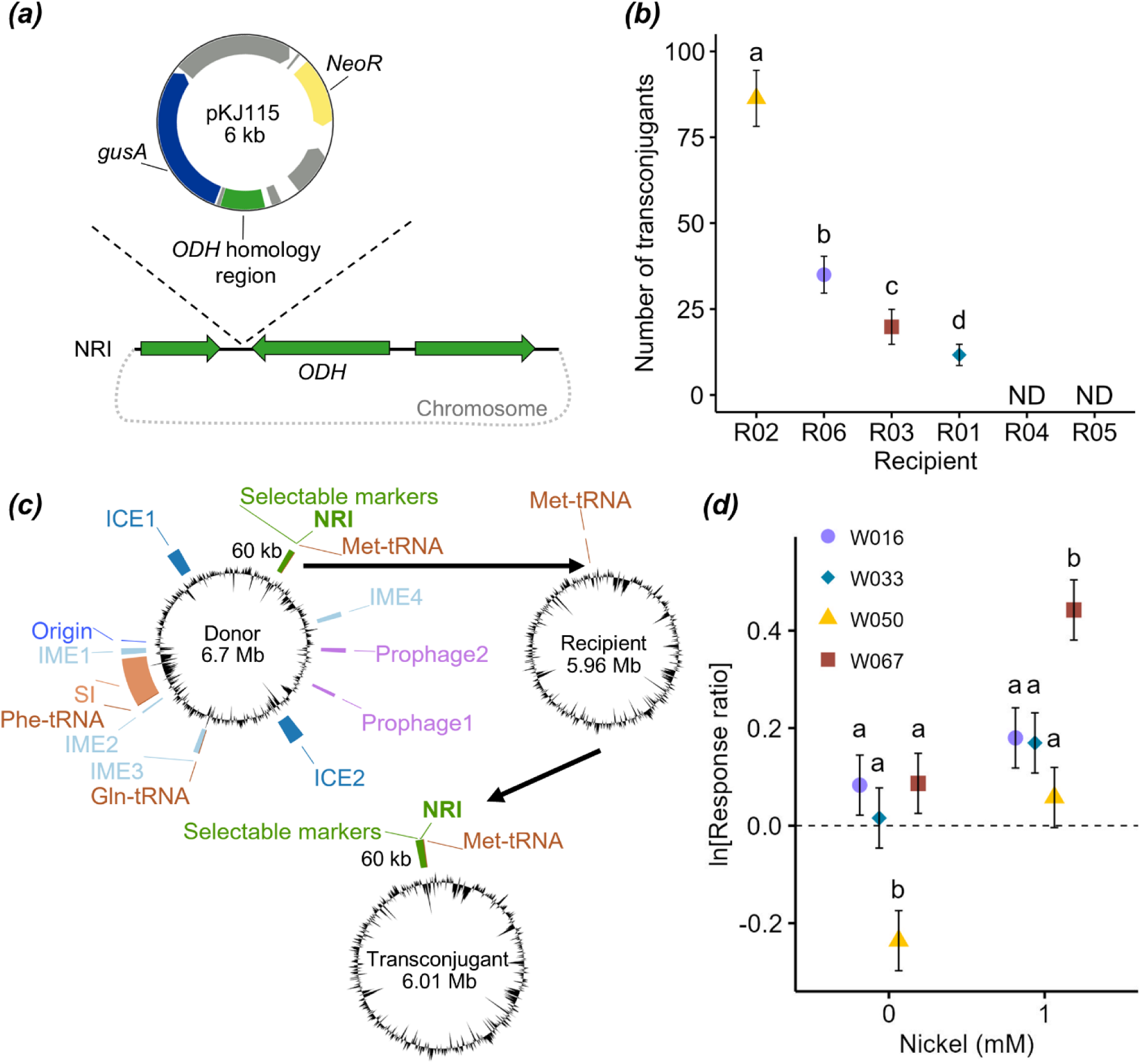
NRI is transmissible *in vitro* where acquisition confers nickel resistance. (a) Insertion of selectable plasmid-borne genes into an NRI intergenic region in donor strain C089B enabled detection of NRI transfer in mating trials (beta-glucuronidase (*gusA*); neomycin resistance (*NeoR*) (see Figure S1 for additional detail). (b) Mean number of transconjugants +/- standard error in pairwise mating trials. Prior to matings, recipients R04 and R06 contain an NRI at the Met-tRNA insertion site while others lack the NRI, but recipients R02 and R05 have putative genomic islands at the Met-tRNA. Four of six recipient *Mesorhizobium* strains accepted the C089B NRI (None detected (ND)) and differed in the number of transconjugants: letters above points indicate significant difference (*P* < 0.05) among strains from an estimated marginal means post hoc test. (c) Example of a mating assay. The donor transferred its marked NRI to recipient R03 which lacked the NRI and is Sm and Rf resistant. In the transconjugant, W067, the NRI inserted at a conserved Met-tRNA gene. The donor bears multiple putative MGEs (Table S2). GC content is shown in black. (d) Log response ratio (ln[RR]) of transconjugant growth in the presence (1 mM) or absence (0 mM) of nickel enriched media. Shown are estimated marginal means. Error bars indicate 95% confidence intervals and those that do not contain zero are considered to have a significant growth response compared to the ancestral strain. Letters above points indicate significant differences (*P* < 0.05) among strains within each nickel level from the estimated marginal means post hoc test.

We find that the donor strain NRI appears to be an integrative and mobilizable element (IME, (45)) capable of excising from and integrating into compatible *Mesorhizobium* chromosomes at a conserved tRNA gene, despite lacking loci for a putative conjugative transfer system. NRI transconjugants were rare, but detectable for 4/6 recipient strains in the mating trials (Table S1; Figure 1b), and transfer to the four recipients resulted in an average of 38 transconjugants per mating. Genome sequencing of 4 transconjugants, one from each successful mating (Table S1), showed that 3 recipients that had initially lacked the NRI gained the donor NRI (transconjugants W033, W050, and W067), which inserted at the conserved Met-tRNA gene in the chromosome (Figure 1c; Figure S2). The transconjugant W033 also acquired another ∼56 kb putative IME from the donor which inserted at a Gln-tRNA (Figure S2b). The recipient R06 initially had an ∼85 kb NRI inserted at the Met-tRNA gene and a second ∼13 kb NRI inserted at an Ala-tRNA gene (Figure S2c). Confirmatory sequencing revealed that this recipient’s respective transconjugant (W016) had its ∼85 kb NRI replaced by the donor’s ∼60 kb NRI at the Met-tRNA gene while the smaller NRI remained intact in the chromosome (Figure S2c).

Given the lack of apparent conjugative transfer genes in the donor NRI itself, we investigated MGEs and cargo genes that could facilitate or constrain NRI transmission. In the donor genome, ICEfinder (46) identified two putative ICEs (ICE1 and ICE2) in addition to the SI that could underly NRI mobilization and conjugative transfer, as well as three putative IMEs (IME1, IME2, and IME4) that could co-transfer with the NRI (Figure 1c; Table S2). However, it did not correctly delineate the SI boundaries and did not identify the NRI or the IME3 that was transferred to transconjugant W033 (Figure 1c; Figure S2b; Table S2). While multiple ICEs may have enabled NRI transfer via conjugation, transconjugant genomic sequences reveal that no ICE co-transferred with the NRI. Thus, the NRI may have co-opted the conjugative transfer function from one or more of these ICEs in a potentially exploitative manner. Genome-wide patterns of variation in GC content often showed lower values for the positions of predicted MGEs we highlight within the main chromosome, enhancing our confidence in our MGE delineation (Figure 1c) (47,48). These additional MGEs on the donor strain bear common adaptive gene repertoires for MGEs (45) including inferred systems for phage defense, an antidefense system for MGE maintenance, and antibiotic efflux genes (see Supplemental Information; Table S3-S8). There are also two putative prophages (Prophage1 and Prophage2; Figure 1c; Table S2). While putative MGE cargo genes could enhance metabolism, defense, or have a complementary role related to symbiosis, they require further study to validate their predicted functions.

To determine whether NRI acquisition conferred tolerance to nickel, we quantified growth in the presence and absence of Ni following Kehlet-Delgado et al. (2024). The growth assay contained the wild-type donor, the NRI-marked donor, 4 recipients and 4 respective transconjugants, and a sterile control. Overall, NRI acquisition tended to enhance transconjugant tolerance to nickel compared to the ancestral recipient strain (OD_600_; χ2 = 233, *P* < 0.001; Table S9; Figure S3) and occurred for three of the four transconjugants (strain*Ni interaction: log response ratio of growth, χ2 = 22.1, *P* < 0.001; Table S1; Table S10; Figure 1d). NRI acquisition in transconjugant W067 caused a 56% increase in growth in Ni, and increased growth in the absence of Ni by 9% and thus showed no cost of NRI carriage (TableS1; Table S10; Figure 1d). Replacement of an existing NRI with the donor NRI in transconjugant W016 increased growth in the presence and absence of Ni by 20% and 9%, respectively (Table S1; Figure 1d). Here the donor NRI appears to have replaced a generally suboptimal NRI variant. NRI acquisition in transconjugant W033 enhanced growth in Ni by 19% with no impact on intrinsic growth rate in the absence of Ni, consistent with gain of a beneficial MGE with no detectable cost to NRI carriage (Table S1; Figure 1d). In contrast, NRI acquisition by transconjugant W050 had no net effect on growth in Ni and incurred a 21% decrease in growth in the absence of Ni (Table S1; Figure 1d; Figure S3). This overall growth reduction is consistent with molecular interference between the donor NRI and this recipient chromosome overshadowing any fitness benefit from the NRI in the presence of nickel. Consistent with long term co-adaptation between the NRI and chromosome attenuating such conflicts between chromosomal- and MGE-encoded molecular machinery, the donor NRI in its native chromosomal background had greater growth in the presence of nickel than any transconjugants (Table S9; Figure S3) (11).

### Imperfect phylogenetic congruence among genomic compartments

To determine whether *Mesorhizobium* genomic compartments are phylogenetically congruent, we used multiple cophylogenetic methods in pairwise tests among the NRI, chromosome, and SI phylogenies. The core chromosomal genome phylogeny was previously constructed for 302 strains using 1,542 single-copy core genes, which are present in every strain (42). The MGE phylogenies were constructed from single-copy core genes in each genomic compartment, but we included multiple copies of *nreA* and *nreX* for 32 strains as these were the only core genes in the NRI (see Methods). The NRI phylogeny was constructed for 208 strains that contain *nreAX* and we included multiple copies, yielding 241 tips on the phylogeny (Figure S4a). 21 single-copy genes were core to the SI (Table S11). The SI phylogeny included 295 strains because 3 strains lacked the SI and 4 strains contained partial SIs that lacked many of the core SI genes present in other strains and were removed. We applied pattern-based (PACo and Mantel) and event-scoring (eMPRess) cophylogenetic methods to test for cophylogenetic signal and phylogenetic congruence, respectively.

We find significant but imperfect cophylogenetic signal and topological congruence between MGE and chromosomal phylogenies. We define the strength of cophylogenetic signal following Perez-Lamarque and Morlon (2024), where low signal is indicated by a PACo R^2^ < 0.25 and a high signal is indicated by PACo R^2^ > 0.5. The chromosome and NRI cophylogeny shows weak cophylogenetic signal (PACo R^2^ = 0.11, *P* < 0.001; Mantel *r* = 0.28, *P* = 0.001; Table S12) though the phylogenies are more concordant than would be expected by chance (eMPRess *P* = 0.009; Figure 2a). Collapsing nodes with low support in the NRI phylogeny resulted in a large polytomy and some well supported groups from which we defined eight phylotypes (N1-N8) (Figure 2a; Figure S4a). The NRI phylogeny aligns broadly with major chromosome clades, for example phylotype N2 is common for clade C1 strains and phylotypes N3 and N4 are common for clade C2 strains (Figure 2a,f).

**Figure 2.**
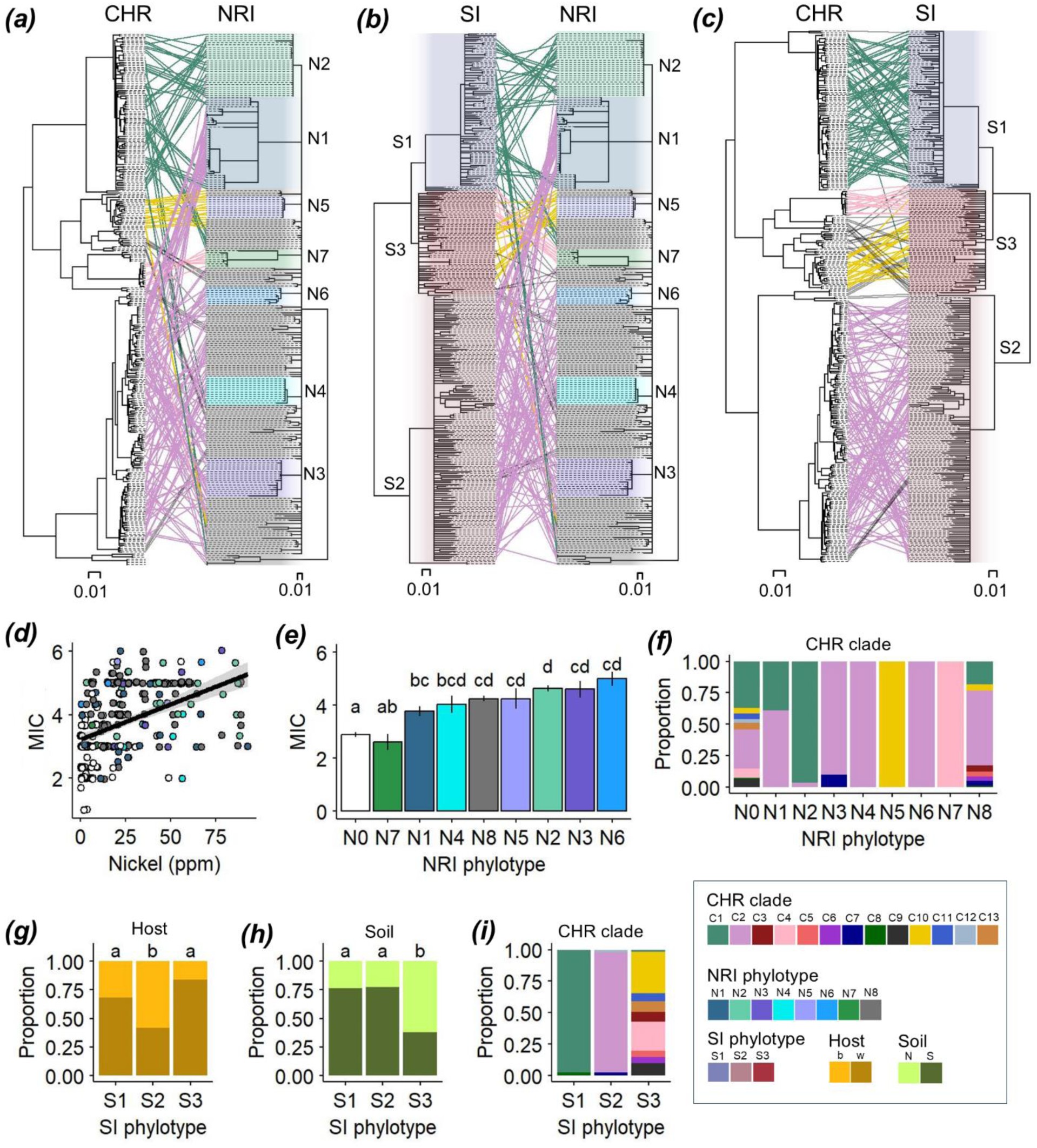
Imperfect phylogenetic congruence between *Mesorhizobium* genomic compartments. Cophylogenies among genomic compartments are more congruent than expected by chance and range between (a) low congruence between the chromosome and NRI, (b) moderate congruence between the SI and NRI, and (c) high congruence for the chromosome and SI. Cophylogenies were built with phytools using NRI, chromosome, and SI phylogenies (RAxML) derived from amino acid sequences of two core genes (*nreA* and *nreX*), 1,542 single-copy core genes, and 21 single-copy core genes respectively. Links between trees are shown for *Mesorhizobium* clades with >10 strains. Nodes with bootstrap support values <70 are collapsed into polytomies. Scale bar indicates the number of amino acid substitutions per site. (d-i) MGE phylotypes assort across abiotic and biotic conditions, consistent with a response to variable selection across environments. (d) MIC (mM nickel) is correlated with soil nickel level (ppm) (*r* = 0.45, *P* < 0.001). Black line; Linear regression. (e) The MIC mean +/- standard error for each NRI phylotype. Bars are colored according to phylotype. N0 indicates the absence of NRI (white). Letters indicate significant differences among phylotypes (*P* < 0.05) based on an estimated marginal means post hoc test. (f) The proportion of chromosomal clades bearing each NRI phylotype. The proportion of (g) host type (*Acmispon brachycarpus* “b”, *A.* wrangelianus “w”), (h) soil type (non-serpentine “N”, serpentine “S”), and (i) chromosomal clade bearing each SI phylotype.

The chromosome and SI cophylogeny shows strong cophylogenetic signal (PACo R^2^ = 0.67, *P* < 0.001; Mantel *r* = 0.92, *P* = 0.001; Table S12) and phylogenetic congruence (eMPRess *P* = 0.009; Figure 2c). The SI phylogeny contains three phylotypes (S1-S3; Figure S5), which largely align with deep chromosome clades (Figure 2c,i). For example, SI phylotypes S1 and S2 have distinct associations with major chromosome clades C1 and C2 strains, respectively, while phylotype S3 aligns with strains from several rare clades (Figure 2b,i).

The SI shows more consistent alignment with the chromosomal phylogeny than does the NRI. For example, chromosomal clade C2 strains only contain SI phylotype S2 (Figure 2c,i), but clade C2 strains can contain diverse NRI phylotypes (Figure 2a,f). Among the MGEs, the SI and NRI cophylogeny (Figure 2b) has moderate cophylogenetic signal (PACo R^2^ = 0.32, *P* = 0.001; Mantel *r* = 0.24, *P* = 0.001; Table S12) but the phylogenies could not be reconciled in eMPRess due to the prevalence of polytomies.

We find no evidence that spatial or geographic barriers to horizontal transmission among *Mesorhizobium* haplotypes cause the cophylogenetic patterns we observe, despite the large spatial extent of our sampling effort. Haplotypes in all three genomic compartments show broad biogeographic distributions across the 1000 km extent of the study (Figure 3) and there is no support for isolation by distance (Mantel r < 0.02 for all compartments; Table S12). For example, well sampled reserves (≥ 18 isolates sampled) contain a diverse mix of haplotypes, with an average of 6 NRI phylotypes, 3 SI phylotypes, and 6 chromosome clades per reserve (Figure 3). Distinct haplotypes also co-occur at spatial scales < 12 m, within individual sites. For example, Hopland site H092c contains three NRI phylotypes (N6, N7, N8), all three SI phylotypes (S1-S3), and four chromosome clades (C1, C2, C3, C4). The NRI has the greatest nucleotide diversity (NRI, 0.065) among the genomic compartments and the chromosome has greater nucleotide diversity than the SI (CHR, 0.048; SI, 0.021; Table S13). The chromosome has slightly reduced GC content compared to the SI and NRI (CHR, 59%; SI, 60.5%; NRI, 62%; Figure 1c; Table S13).

**Figure 3.**
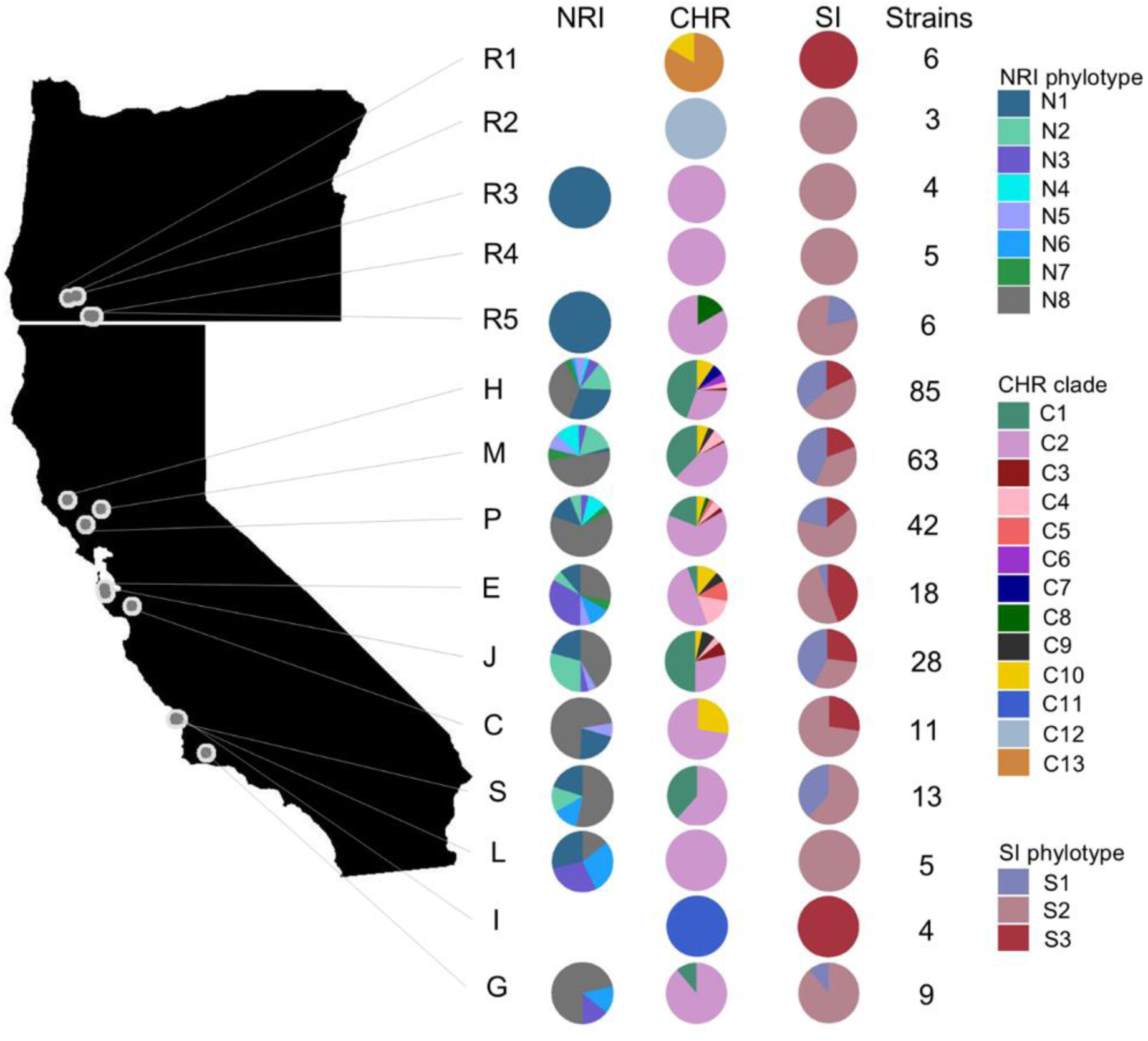
Widespread co-occurrence of haplotypes among and within reserves. Shown are the proportion of strains within each reserve from different NRI phylotypes (N1-N8), chromosomal clades (C1-C13), and SI phylotypes (S1-S3). The number of strains per reserve is shown. We include strains with multiple copies of NRI core genes. Reserves include: R1, Oregon 1; R2, Oregon 2; R3, Oregon 3; R4, Oregon 4; R5, Oregon 5; H, Hopland Research and Extension Center; M, McLaughlin Natural Reserve; P, Pepperwood Preserve; E, Edgewood Park & Natural Preserve; J, Stanford University Jasper Ridge Biological Preserve; C, Kirby Canyon Landfill; S, South Hills Open Space; L, Laguna Lake Park; I, Irish Hills Natural Reserve; G, Sedgwick Reserve.

### MGE phylotypes confer distinct phenotypes

Allelic variation (i.e., differences among phylotypes) at the NRI and SI contributes genetic variation that has the potential to fine-tune environmental adaptation. We tested for ecological differences among phylotypes in the NRI or SI and among clades for the chromosome using multivariate regression and Fishers exact tests (see Methods). Consistent with their adaptive significance, NRI phylotypes are associated with differences in nickel tolerance (MIC) and the level of Ni enrichment in origin soil, and SI phylotypes differ in the frequency with which they associate with the two host plant species. The NRI phylotypes we delineate are associated with different nickel MIC and soil nickel level (Wilks’ lambda = 0.72, F_16, 506_ = 5.0, *P* < 0.001; Figure 2d,e). Furthermore, NRI phylotypes that confer greater nickel tolerance tend to originate in soils with higher levels of nickel (Figure 2d,e). In contrast, SI phylotype did not predict MIC and soil nickel level. The association between chromosomal clade and MIC and soil nickel level (Wilks’ lambda = 0.78, F_24, 562_ = 3.0, *P* < 0.001) appears to be driven by clades that were only observed on non-serpentine soil (Figure S6). Similarly, some SI phylotypes are more common on a particular soil type. In particular, SI phylotype S3 is more common on non-serpentine soil than phylotypes S1 and S2 (Fisher Test *P* < 0.001; Figure 2h).

SI phylotypes and chromosomal clades are differentially associated with the two host species. SI phylotypes S1 and S3 are prevalent in strains from host *A. wrangelianus* while SI phylotype S2 are more common in strains from *A. brachycarpus* (Fisher Test *P* < 0.001; Figure 2g). The chromosome’s phylogenetic congruence with the SI recapitulates patterns with host species, where chromosome clade C1 (S1 associated) and C10 (S3 associated) are more common with host *A. wrangelianus* while clade C2 (S2 associated) is more common with the host *A. brachycarpus* (Fisher Test *P* < 0.001; Figure S6). NRI phylotypes are also differentially associated with the two host species (Fisher Test *P* = 0.008), though this effect may be driven by rare observations of some phylotypes (Figure S4b).

### NRI loss accompanies migration to non-serpentine soils

We hypothesized Ni adaptation evolved under conditional neutrality because the NRI confers Ni tolerance and is nearly ubiquitous in strains from Ni-enriched serpentine soils, but incurs no detectable cost of carriage in the absence of Ni and is largely absent in strains from non-serpentine soil (42,49). To investigate evolutionary dynamics under apparent conditional neutrality, we used ancestral state reconstruction with discrete traits across the phylogeny of *Mesorhizobium* in our focal population to investigate whether inferred *Mesorhizobium* migration across soil types corresponded to NRI evolutionary gain or loss. Ancestral state reconstruction predicted that the most common evolutionary transition was for ancestral lineages that inhabit serpentine soil and bear the NRI to migrate to non-serpentine soil and lose the NRI. This transition in trait states was predicted to be more common than for all other transitions among possible states (Figure 4b; Table S14), including for the reverse transition, wherein ancestral lineages that inhabit non-serpentine soil and lack the NRI migrate to serpentine soil and gain the NRI, which has overlapping HPD intervals with several other transitions (Table S14). Thus, migration of strains from serpentine to non-serpentine soil concomitant with NRI loss appears to have been common in the evolutionary history of the focal *Mesorhizobium* bacteria (Figure 4; Figure S7).

**Figure 4.**
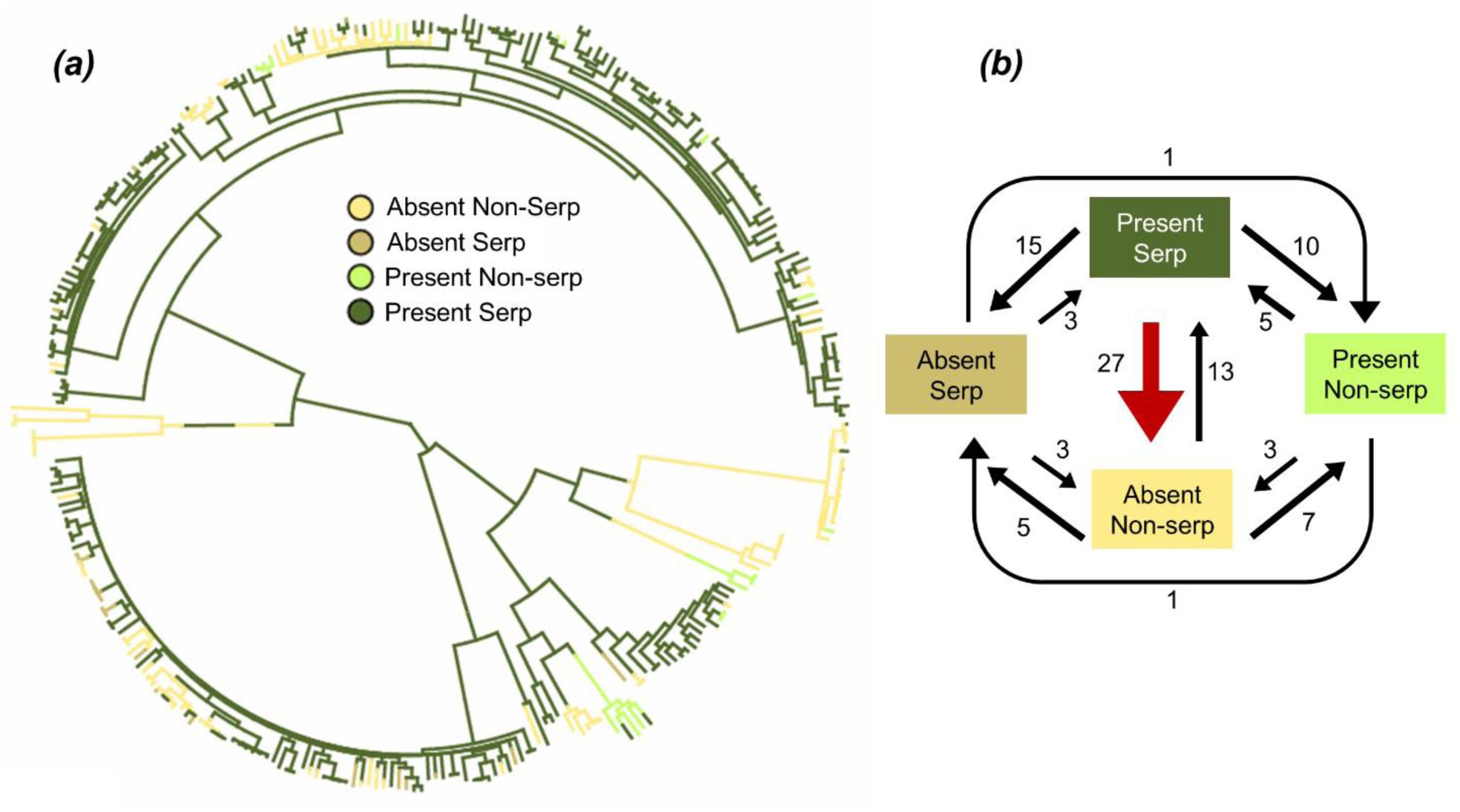
Ancestral trait reconstruction indicates frequent NRI loss upon migration from serpentine to non-serpentine soils. (a) Ancestral state reconstruction and (b) transition path diagram for NRI character states based on the presence or absence of NRI on serpentine or non-serpentine soils. Shown is one stochastic character map from the sampled character histories consistent with tip data (see Figure S7 for posterior probabilities of 1000 stochastic maps). Tip color indicates the known state for each taxon. Branch colors indicate the predicted evolution of the states under the equal transition rates model. Arrow size is proportional to estimated transitions rates (see Table S14 for high probability density (HPD) values for each transition).

## DISCUSSION

The bacterial mobilome encompasses a vast reservoir of heritable variation that often underlies local adaptation. Yet, contemporary and ancient transmission dynamics for adaptive MGEs remain largely unexplored in natural landscapes (7). Transmission of adaptive MGEs is shaped both by natural selection for tolerance to environmental conditions and by genetic and spatial constraints on transfer. The balance between these factors could drive scenarios that range from phylogenetic congruence to independence between MGEs and the host chromosome they reside in. We demonstrate the horizontal transmissibility of an MGE and its adaptive function for heavy metal tolerance and reveal an intermediate scenario in which wild *Mesorhizobium* show imperfect phylogenetic congruence among adaptive MGEs and the recipient genome.

Phylogenetic congruence of the NRI, SI, and chromosome is consistent with frequent vertical co-inheritance and/or MGE transmission primarily among close relatives over deep timescales (17,21,27,50). This congruence persists even without apparent spatial barriers to MGE transmission: MGEs and chromosome haplotypes have broad biogeographic distributions and co-occur at fine spatial scales. The evolutionary rarity of MGE transmission among distant relatives is consistent with strong selection to maintain co-adapted genomic compartments (51,52). When adaptive MGE loss does occur, it often appears to coincide with migration across environments. Here, spatial variation in selection due to heterogenous environmental factors such as soil type, may occasionally be sufficient to disrupt phylogenetic congruence and co-adapted gene complexes among genomic compartments and serve as an engine of diversity generation by favoring novel chromosomal-MGE haplotype combinations as strains adapt to novel conditions (23,25).

### MGE transmission dynamics over short timescales

Understanding how MGE transmission potentiates yet constrains wild bacterial local adaptation is a critical frontier with broad implications for the role of MGEs in expanding or contracting microbial niches (8). The NRI enables wild *Mesorhizobium* to adapt to stressful nickel-enriched serpentine soils and drives geographically widespread and replicated nickel-adaptation across natural landscapes (42). Horizontal transmission of the NRI was previously inferred due to imperfect linkage between NRI and chromosomal alleles (42). We provide molecular confirmation that an NRI variant is transmissible, despite lacking putative genes for conjugative transfer, consistent with its classification as an integrative and mobilizable element (IME) (1). Transfer of this NRI into a conserved Met-tRNA in the chromosome of recipient strains is detectable but rare, and may rely upon one or more integrative and conjugative elements (ICEs) in the donor chromosome for conjugative transfer (53). This NRI variant is capable of acquisition by both recipient strains that had previously lacked the NRI, and by a strain that had already carried two distinct copies of the NRI, one of which it replaced.

For facultative endosymbionts like rhizobia, the genetics and genomics of adaptation to niches outside of the host plant have been challenging to elucidate (54,55). Our experimental NRI transmissions reveal that strong chromosome-MGE epistasis shapes environmental MGE niche-specific adaptation of rhizobia to soil habitats: both the number of transconjugants as well as functional consequences of NRI acquisition differed among recipient strains. Only four of six tested recipient strains successfully acquired the NRI, and these strains varied 7-fold in the number of transconjugants produced during mating experiments. While this NRI variant increased nickel tolerance in recipient strains overall, recipient strains also varied 7-fold in the magnitude of its effect on nickel tolerance, and 3-fold in its overall carriage costs.

The NRI appears to have the potential to evolve under dynamics that approach conditional neutrality (56), as it enhances growth in the presence of nickel, but does not detectably inhibit growth in nickel’s absence for many strains *in vitro.* Expression of putative efflux pumps encoded by *nreX* and *nreY* is induced by *nreA,* a transcriptional repressor responsive to exogenous Ni that binds a promoter immediately upstream of the *nreAXY* operon (44). Thus, the *nre* operon has both Ni-inducible expression and a compact size and organization. This may predispose this operon to incur only weak carriage costs (57), consistent with our experimental findings which did not reveal an impact of NRI gain on growth *in vitro* in the absence of Ni for most strains. Despite this apparent conditional neutrality, the absence of NRI in strains from non-serpentine soils, where nickel tolerance is not advantageous, could still evolve due to mutation accumulation and loss of the MGE under relaxed selection where Ni is not present in the environment (49).

However, weak carriage costs would have gone undetected in our growth assays and could still lead to MGE-chromosomal co-adaptation over many generations, especially in chromosomal backgrounds where carriage costs are higher. The *nre* operon or other less well-studied NRI loci might interfere host chromosome gene networks in other environmental or genomic contexts. Reflecting this potential for MGE-chromosome genetic incompatibility, NRI acquisition imposed a high carriage cost for one recipient strain such that it hampered growth in the absence of nickel enrichment and did not significantly enhance growth in the presence of nickel. Furthermore, we found that the NRI donor strain had greater growth in the presence of nickel than any transconjugants that received its NRI. This is consistent with a scenario in which nickel tolerance can be optimized when an NRI is in its native chromosomal background. Here, co-adapted gene complexes and compensatory mutations may accumulate over evolutionary time to reduce gene network conflicts, even if they are weak (11). Such intra-genomic co-adaptation could, over time, create genomes comprising multiple replicons that together provide high fitness, but the resulting co-dependence may limit the spread of co-adapted MGEs to other lineages (58).

### Deep phylogenetic congruence consistent with MGE-chromosome co-adaptation

Consistent with selection that favors co-adapted MGE-chromosome combinations, we find remarkable levels of cophylogenetic signal and phylogenetic congruence among the NRI, SI and chromosome in wild *Mesorhizobium* genomes. This congruence is maintained despite the opportunity for horizontal transmission among co-occurring MGE and chromosome haplotypes which show broad biogeographic distributions across the 1000 km extent of this study and co-occur within individual sites (12 m). The NRI shows evolutionary congruence with the chromosome, but at lower levels than for the SI. This is consistent with the NRI experiencing less intragenomic epistasis and gene network interaction complexity than the SI (42). The NRI had low cophylogenetic signal with the chromosome and moderate cophylogenetic signal with the SI. Despite the overall signal of phylogenetic congruence between the NRI and chromosome, cophylogeny links indicate that core chromosome clades and 7 NRI phylotypes are imperfectly aligned (59). This incongruence is also evident in the NRI and SI cophylogeny. Cophylogenetic signal between these two MGEs appears greater than that between the NRI and chromosome, though this could arise due to the relatively low diversity of SI phylotypes. The stronger codependence between the chromosome and SI compared to NRI is similar to other studies in rhizobia that find stronger patterns of codependence between genomic compartments that carry complementary genes important for survival or transfer (60–62).

The evolutionary patterns of MGE transmission across natural landscapes we observe are concordant with theoretical predictions based on intra-genomic molecular interactions. The complexity hypothesis posits that higher levels of gene network complexity in an MGE result in stronger intragenomic epistatic constraints on the phylogenetic breadth of MGE transmission (3,14). Symbiosis MGEs like the SI encode proteins that underlie complex molecular interactions with proteins encoded by both the recipient genome and the plant host genome. For example, symbiosis gene expression on plasmid pSymA in *Sinorhizobium meliloti* is responsive to both chromosomal and *Medicago truncatula* host plant gene coexpression (37). The SI is also four-fold larger than the NRI and thus has a higher potential for protein interaction network complexity than does the NRI. In contrast, for stress tolerance MGEs like the NRI, tolerance is often conferred by a few inducibly expressed genes, which are predicted to have lower gene network complexity and interaction with the main replicon. Concordant with evolutionary outcomes that could be predicted from the complexity hypothesis, we find higher cophylogenetic signal and topological congruence with the chromosome for the SI than for the NRI (27,59).

Over time, these greater differences in complexity could lead to greater co-adaptation between the SI and chromosome than for the NRI and chromosome, such that co-adapted gene complexes are a stronger constraint on the phylogenetic breadth of SI transmission (58).

Our findings highlight the importance of co-adaptation between MGEs and recipient genomes in shaping diversity in rhizobia, an important group of plant symbionts responsible for roughly half of all terrestrial nitrogen fixation (63). Phylogenetic congruence is apparent in patterns such as the fact that SI phylotype S1 resides within chromosomal clade C1, S2 resides within chromosomal clade C2, and S3 resides in several minor clades, with few exceptions.

These types of patterns in our study are especially strong compared with levels of cophylogenetic signal between the chromosome and symbiosis gene elements observed in other rhizobium pangenome studies, which is highly variable among studies (61,62,64,65). For example, Wang et al. (2018) observed moderate cophylogenetic signal between the chromosome and symbiosis gene elements among several rhizobium genera but did not find phylogenetic congruence, while Karasev et al. (2023) found perfect phylogenetic congruence among two biovars of *Neorhizobium galegae*. In contrast, Hollowell et al. (2016) found no cophylogenetic signal between *Bradyrhizobium* chromosomes and symbiosis islands across a scale of sampling of wild rhizobia similar to the present study. Our findings are in contrast with those from some studies that infer phylogenetic incongruence by visually inspecting chromosome and symbiosis gene element phylogenies for evidence of HGT (66,67), though these studies do not tend to test the inference statistically. It is possible that incomplete phylogenetic resolution in previous studies could have obscured phylogenetic congruence, as phylogenies are often inferred from relatively few core genes, rather than all single-copy core genes as in the current study (n= 1542 for chromosome, n = 21 for SI). Variation among published studies in the strength of MGE-chromosome cophylogenetic signal and phylogenetic congruence may also simply reflect substantial variation among rhizobia populations in the fidelity with which symbiosis MGEs are inherited and transmitted horizontally.

### Mosaics of selection and imperfect congruence

The cophylogenetic signal and congruence we observe are imperfect. Environmentally variable selection on MGE-conferred function could contribute to deviations from cophylogeny among genomic compartments. As *Mesorhizobium* lineages migrate across environments, there may be spatially variable selection that selects for different MGE variants in different locations. In the case of the NRI, different phylotype variants confer distinct levels of tolerance to nickel, which is enriched to varying degrees at different serpentine sites (42). Thus, selection on different serpentine soil sites for nickel tolerance may favor acquisition of distinct locally matched NRI phylotypes by chromosomal lineages that colonize the sites. This potential for evolutionary fine-tuning of MGE-conferred local adaptation contrasts with other systems in which MGE variants don’t show detectable differences in their impact on function. For example, subtypes of the mobilizable element TR1 in *Streptomyces*, a pathogen in potato, do not show detectible differences in virulence (68), and thus may experience microevolutionary processes that are distinct from those we observe.

Heterogeneously distributed host plant species can also impose spatially variable selection that could favor different SI haplotypes in different locations. Most of the genes required for intimate symbiosis with a host plant reside in the SI (5,69,70) and different hosts can select for different SI variants, such as distinct nodulation factor alleles (62). Correspondingly, we find that *Mesorhizobium* SI phylotypes are differentially associated with the two host species we examined. SI phylotypes S1 and S3 are more common on *Acmispon wrangelianus,* and SI phylotype S2 is more common on *A. brachycarpus,* despite the fact that these host species often co-occur at sites. Finding host-associated phylotypes is consistent with other studies that find associations between variants of symbiosis gene elements and different host species for other rhizobia populations (38,60,62). Host plant species also differentially associate with some chromosomal clades, which could be due either to phylogenetic tracking between the chromosome and SI, or due to selection for particular chromosome clades on a host species.

Our findings implicate both soil chemistry and legume host distributions as factors shaping the spatial distribution of *Mesorhizobium* pangenome adaptive diversity. Over generations, as chromosomal lineages migrate across environments, spatially variable selection for different adaptive MGE variants may periodically disrupt co-adapted MGE and chromosome combinations by favoring a locally adaptive MGE variant. In this way MGE-conferred local adaptation could serve as an engine of diversity by periodically favoring novel MGE-chromosomal associations as lineages migrate across environments.

### Mosaics of selection and MGE gains and losses

Ancestral state reconstructions across the *Mesorhizobium* phylogeny indicate that NRI loss is associated with a predicted ancestral migration from serpentine to non-serpentine soil. This is concordant with other studies that observe MGE loss in the absence of environmental stress (3,18,71). However, the NRI often shows approximately conditionally neutral fitness effects *in vitro.* If the NRI confers only a weak cost in the absence of nickel, its loss could result due to relaxed selection combined with bacterial genomic streamlining in non-serpentine soil (3). Ancestral trait reconstructions also suggest that the loss of the NRI upon migration from serpentine to non-serpentine soil habitats occurs more frequently than the reverse, gain of the NRI upon migration from non-serpentine to serpentine soil. This is consistent with a scenario in which lineages that migrate to non-serpentine soil may persist long enough to eventually lose the NRI, whereas lineages that lack the NRI and migrate to serpentine soil tend to go extinct before they gain the adaptive NRI. Even if a strain acquires the NRI on serpentine soil, our transmission experiments indicate that it will typically not confer full function immediately or could even impose a cost, as observed in transconjugant W050, which could lead to extinction of lineages that have migrated to serpentine soil. Future investigations on asymmetric patterns of MGE gain and loss upon migration between environments would be valuable for bolstering evidence for such polarized evolutionary dynamics.

The high phylogenetic congruence between SI phylotypes and chromosomal clades we observe indicates that the SI is primarily vertically inherited and/or horizontally exchanged among close relatives. High rates of vertical transmission for the SI is consistent with previous findings wherein SI loss is very rare *in vitro* and symbiotic mutualism is robust to instability due to the shared and substantial fitness benefits for plants and rhizobia (29,72–74). MGEs often evolve molecular strategies that prevent their loss during cellular replication, such as toxin-antitoxin systems, which occur in *Mesorhizobium* SIs and other rhizobium symbiosis MGEs (39,75,76). Furthermore, rhizobia symbiotic elements are not known to impose large costs of carriage, though they may divert small amounts of resources from growth or competitive ability in the absence of hosts (77). In the root nodule isolates we examined, the SI is ubiquitous and present in 98% of strains, in contrast to the NRI, which is absent from 31% of strains.

Cophylogenetic patterns are consistent with restriction of most horizontal SI transmission to lineages from the same ancient chromosomal clades. This pattern is consistent with negative epistatic effects arising due to co-adapted gene complexes that restrict the SI to vertical transmission or horizontal transmission among closely related relatives. These findings correspond with epistatic constraints identified in other systems, where transfer of symbiotic elements into different rhizobia chromosomal backgrounds can lead to a failure to form nodules or fix nitrogen (34,78), though these restrictions are not ubiquitous (32).

We characterize the NRI in reference strain C089B as an IME. However, the NRI appears to bear molecular features that span other classes of MGEs, possibly along the continuum of MGE ‘domestication’ (8,79). Based on homology, NRI organization as an IME appears common in the focal *Mesorhizobium* population (42). However, in some strains the NRI bears features consistent with an ICE such as putative conjugative type IV secretion systems, and in other strains it appears to comprise a circular replicon consistent with a plasmid (42). Future research will be required to understand how conversion from one form into another evolves for adaptive MGEs, why the adaptive loci from these MGEs are not more frequently domesticated into the main replicon, and the implications for bacterial local adaptation.

## CONCLUSION

We reveal how tensions between environmental selection, transmission opportunities, and haplotype stability shape evolutionary trajectories for distinct compartments within bacterial pangenomes. While MGEs that confer locally adaptive traits can speed bacterial adaptation, the spread of MGEs is shaped by multiple constraints, which result in distinct evolutionary trajectories from their host genome (8). Adaptive MGEs and the replicons they reside in may face tradeoffs as the phylogenetic breadth of transmission evolves. On one hand, adaptations that optimize fitness in a particular chromosomal background could lead to specialization of transmission among a phylogenetically narrow set of chromosomal diversity, as the SI appears to do. On the other hand, an MGE could acquire more generalizable adaptations that enable successful transmission across a broader set of chromosomal diversity, enabling more generalized horizontal transmission, as the NRI in this study appears to do (3,16,80,81). Further integrative research to connect adaptive loci and traits to their fitness effects on the multiple replicons within bacterial cells will be critical to shed light on how bacteria adapt to heterogeneous natural landscapes.

## METHODS

### NRI transmission and function in vitro

The donor NRI from strain C089B is an IME that lacks conjugation genes (42), but could be mobilized by an MGE that contains conjugation machinery, such as the SI. SI transmission is inducible and transmission rates can increase in the presence of host plants (82,83). Therefore, matings were performed using plant root extracts that can stimulate SI transfer (82,83). Insertion of the marker plasmid into the donor’s NRI showed no detectable effect on growth of the donor strain in the presence or absence of nickel (Table S9; Figure S3). The donor is neomycin (Nm) resistant and blue in the presence of X-Gluc, while recipient cells were selected for streptomycin (Sm) and rifampicin (Rf) resistance and are colorless in the presence of X-Gluc. Novel transconjugants that result from the transmission of the NRI from donor to recipient are Nm, Sm, and Rf resistant and blue in the presence of X-Gluc. Strains were incubated at 28°C on Tryptone Yeast (TY) agar (84) and as appropriate supplemented with 100 μg/mL Nm, 200 μg/mL Sm, 100 μg/mL Rf, and/or 50 μg/mL X-Gluc.

In pairwise mating trials, NRI donor and recipient cells from plates were suspended in TY broth and normalized to 6 × 10^8^ cells per mL using the formula, cells mL^−1^ = OD_600_ × 5.8 × 10^7^ (85). 30 μL mating spots with equal ratios of donor and recipient cells and 10 μL of plant root extracts, were incubated on TY agar for 40 hours. Entire mating spots were gathered with a sterile pestle, suspended in 350 μL of TY broth, and 100 μL was spread on three replicate selection plates to count colony forming units (CFUs) to quantify the number of transconjugants per mating. Negative controls, consisting of only donor or recipient cells, mixed with plant root extracts, were also plated on selection plates. Colonies from selection plates were transferred to TY agar supplemented with X-Gluc and Nm, reisolated as single colonies, cultured, and cryopreserved. We tested whether recipient strains that gained the NRI differed in the number of transconjugants using a linear model.

Strains used in the growth assay (Table S1) were genome sequenced by Plasmidsaurus using Oxford Nanopore Technology with custom analysis and annotation and compared to genomic sequences from Kehlet-Delgado et al. (2024) to: 1) confirm integration of the plasmid marker into the donor NRI, 2) confirm NRI acquisition in transconjugants, and 3) confirm the identity of the chromosome in transconjugants. Sequence alignments were performed between the wild-type donor and NRI-marked donor or respective recipient-transconjugant pairs using *progressiveMauve* (86), and genomic maps were visualized in proksee (87). To check whether recipients that lack the NRI at the Met-tRNA gene contain other putative genomic islands (GIs) at this insertion site, we used IslandViewer4 (88) which predicts GIs using multiple methods based on sequence composition and comparative genomics. Predicted GIs were evaluated further by identifying direct repeats of the end of the Met-tRNA gene and examining whether this region aligned with a divergence in GC content.

To identify MGEs with conjugation machinery that could mobilize the NRI within the donor strain, we used ICEfinder to identify MGEs based on annotated features, such as integrase genes or type IV secretion systems (46). We explored gene content within each MGE predicted by ICEfinder and an additional IME identified informatically during the transmission experiment. We tested for functional enrichment in putative MGEs compared to the background genome of the donor with FUNAGE-Pro (89). To investigate whether putative MGEs carry cargo genes associated with resistance to phage or antibiotics we examined them with Defense-Finder (v2.0.0) (90,91) and CARD (v4.0.0) Resistance Gene Identifier (RGI) (v6.0.3) (92), respectively. Putative prophage were identified using Phaster (93) and geNomad (94).

To determine whether NRI acquisition conferred tolerance to nickel, we measured strain growth in the presence or absence of Ni following Kehlet-Delgado et al. (2024). Briefly, we resuspended cultured strains at 1 × 10^6^ CFU/mL (based on OD_600_), inoculated 5 μL into a 384-well plate incubated in a TECAN microplate reader at 28°C, and measured growth (OD_600_) at 72 hours. The assay manipulated both Ni level (in 0 or 1 mM NiCl_2_), and strain (wild-type donor, the NRI-marked donor, 4 recipients and 4 respective transconjugants, and a negative sterile control) in a complete randomized block design with 5 blocks. To test how NRI acquisition impacts growth, we used the log response ratio (ln[transconjugant growth/recipient growth]) as the response variable in a GLMM (95) with an interaction of Ni level and strain as fixed effects and block as a random effect. To check whether the plasmid impacted growth of the donor we also modeled all strains growth predicted by the interaction of strain and nickel level as a fixed effect and block as a random effect. For all models described above, a likelihood ratio test was used to test fixed effects, followed by an estimated marginal means (96,97) post hoc analysis to assess differences among groups. Error distributions were checked using simulated residuals in DHARMa (v0.3.3) (98). Statistical analyses were conducted in R studio (v4.2.2) (99).

### Phylogenetics

The focal *Mesorhizobium* pangenome was previously characterized and the core chromosomal genome phylogeny was constructed for all 302 sequenced strains (286 draft genomes via Illumina and 16 complete genomes via PacBio) (Kehlet-Delgado et al. 2024). Given the nature of the draft genomes, we leveraged the PacBio genomes to identify genes present in the complete NRI and SI. Then, from the *Mesorhizobium* pangenome we identified genes that are present in all strains that contain an NRI or an SI, and excluded genes present in strains that do not contain an NRI or SI. Since we could not differentiate genes within the MGEs from similar gene hits on the main chromosome in the draft genomes due to contig breaks, we focused on single-copy core genes in the MGEs. Only nickel tolerance genes, *nreA* and *nreX,* were core to the NRI, which reflects the high diversity of these genetic elements (42). However, 32 strains had more than one copy of one or both genes within the NRI (42). Since we’ve previously identified these genes and differentiated them from chromosomal homologs in non-serpentine strains (42), we relaxed the single-copy gene requirement for core genes in the NRI. Concatenated amino acid sequence alignments of NRI and of SI core genes were used to build maximum likelihood phylogenies for both MGEs using the “automatic” option in RAxML (100) to determine the best model. JTT amino acid and DUMMY2 substitution models with empirical base frequencies were used for the NRI and SI phylogenies, respectively. Phylogenies were processed in R studio using ape (v5.7.1) (101) to midpoint root and collapse nodes with bootstrap support values less than 70 into polytomies, and visualized with ggtree (v3.6.2) (102).

To define NRI and SI phylotypes, we attempted to calculate pairwise ANI distance between the concatenated alignments of core genes. Due to high levels of divergence, FastANI was unable to estimate all pairwise ANI comparisons for the NRI, so we define NRI phylotypes based on well supported nodes on the amino acid phylogeny (103). However, nodes containing less than eight taxa were binned into NRI phylotype N8. Strains with multiple copies of *nreAX* are included in downstream analyses that permit multiple states, as noted below. We defined SI phylotypes using 97% ANI because this generated clusters that corresponded to well supported nodes on the phylogeny, whereas a threshold of 95% ANI for the SI did not result in distinct clusters (FigureS5a) (103). *Mesorhizobium* core chromosomal diversity comprises two major clades abundant across the landscape, Clade C1 (n = 89) and Clade C2 (n = 144), and 11 minor rare clades (42).

### Cophylogenetic analyses

To test for phylogenetic congruence, we used multiple pairwise cophylogenetic tests. The pattern-based methods we used test whether two phylogenies are more similar than expected by chance. We applied the global-fit test of Procrustean Approach to Cophylogeny (PACo) in R (104,105) on phylogenetic distance matrices of the NRI, chromosome, and SI. We assumed a *symmetric* fit because it is unknown whether one phylogeny tracks the other: the MGEs could track the chromosome phylogeny similar to vertically transmitted symbionts on hosts (21), they could track one another due to collaboration (106), or they could have independent evolutionary trajectories. We used 1000 permutations to refit cophylogenetic links using the “swap” randomization method. Stronger cophylogenetic signal is indicated by a low residual sum of squares (*m^2^_xy_*) from PACo (104,105), and the effect size of cophylogenetic signal in the *symmetric* test, R^2^ (1-*m^2^_xy_*), ranges from 0 (low cophylogenetic signal) to 1 (high cophylogenetic signal) (27,50).

The event-scoring method we used reconciled the discordance between two phylogenies by fitting events, such as duplication, loss, and transfer that could have produced their cophylogenetic structure (20). We used eMPRess (59), to test whether genomic compartment phylogenies were more topologically congruent than expected by chance using maximum parsimony to reconcile the phylogenies by assuming one tracks the evolution of the other. First, we assumed the MGE phylogenies track the chromosome phylogeny. We implemented default event costs to maintain an equal ratio between duplication, transfer, and loss (cost = 1), and considered codivergence a null event (cost = 0). Next, since it’s unclear whether the SI or NRI track the evolution of one another, we attempted reconciliation both ways (i.e., NRI tracks SI or SI tracks NRI). The test for phylogenetic congruence compares the maximum parsimony cost of 100 cophylogenies with random tip associations to the original cophylogeny tip associations.

Therefore, the null hypothesis that two phylogenies are congruent by chance can be rejected at a p-value threshold of 0.01 (59).

We used Mantel tests in R (vegan v2.6-4) (107) to investigate whether among-strain genetic distance matrices correlate with one another between genomic compartments or across geographic distance. The concatenated amino acid sequences of core genes for each genomic compartment were used to create distance matrices using the dismat function in EMBOSS (v 6.6.0) (108) and the Kimura protein distance was used to correct the observed substitution rate. We compared distance matrices between genomic compartments in pairwise tests to investigate the extent to which the background genome influences transfer or relatedness of the MGEs. To test whether the diversity and distribution of the genomic components is geographically structured, we compared genomic compartment distance matrices to a spatial distance matrix based on collection site coordinates. Strains with more than one copy of *nreAX* were excluded from NRI associated mantel tests. As MGEs often show divergent sequence composition, we estimated and compared GC content and nucleotide diversity within the core gene sequences from each genomic compartment.

### MGE-conferred adaptations among phylotypes

To test for differences in the proportion of each phylotype or clade on a particular host species or soil type, we used Fisher’s Exact Test and pairwise comparisons with a Bonferroni correction for multiple tests (rstatix v0.7.2) (109). We also tested whether phylotypes or clades differed in nickel tolerance (MIC) and nickel level in the soil of origin. Environmental soil nickel level predicts *Mesorhizobium* nickel tolerance *in vitro* (42). Therefore, we used multivariate linear regression with the response of MIC and nickel level predicted by MGE phylotype or chromosomal clade and we included soil type (serpentine or non-serpentine) as a control covariate. To improve model fit to assumptions, we used a log transformation for soil nickel level in each multivariate model. MANOVA was followed with an estimated marginal means post hoc test. In addition, we modeled MIC predicted by NRI phylotype in a univariate model to test whether phylotypes confer different levels of nickel tolerance. Strains with multiple copies of *nreAX* were excluded from NRI tests.

### NRI ancestral state reconstruction

We used discrete NRI presence/absence states to infer ancestral trait transitions for strains from serpentine or non-serpentine soil, for a total of four states in stochastic character mapping (phytools v2.1-1) (110). Compared to assumptions of symmetric transition rates, or that all transition rates differ, the assumption of equal transition rates was best supported using Akaike information criterion (AIC) and was used in *simmap* with 1000 simulations. To compare transition rates between NRI states we examine the Bayesian 95% high probability density (HPD) interval using the posterior distribution of sampled character histories (110).

## DATA AVAILABILITY

Genome sequence data have been deposited in NCBI bioproject PRJNA1334436. Data is deposited in the Dryad data repository but is currently private for peer review (DOI: 10.5061/dryad.rv15dv4m4).

## ACKNOWLEDGEMENTS

SSP and APM were supported by US National Science Foundation grant DEB-1943239 to SSP. APM was also supported by the American Association of University Women (AAUW) American Dissertation Fellowship and the Washington State University, School of Biological Sciences, Rexford Daubenmire Fund for Graduate Education. This research used resources of the Center for Institutional Research Computing at Washington State University.

## REFERENCES

1. Lang AS, Buchan A, Burrus V. Interactions and evolutionary relationships among bacterial mobile genetic elements. Nat Rev Microbiol. 2025 Mar 11;1–16.

2. Weisberg AJ, Chang JH. Mobile Genetic Element Flexibility as an Underlying Principle to Bacterial Evolution. Annual Review of Microbiology. 2023 Sep 15;77(Volume 77, 2023):603–24.

3. Brockhurst MA, Harrison E, Hall JPJ, Richards T, McNally A, MacLean C. The Ecology and Evolution of Pangenomes. Current Biology. 2019 Oct 21;29(20):R1094–103.

4. Hall JPJ, Harrison E, Baltrus DA. Introduction: the secret lives of microbial mobile genetic elements. Philos Trans R Soc Lond B Biol Sci. 2021;377(1842):20200460.

5. Heath KD, Batstone RT, Cerón Romero M, McMullen JG. MGEs as the MVPs of Partner Quality Variation in Legume-Rhizobium Symbiosis. mBio. 2022 Jun 27;13(4):e00888–22.

6. Vale FF, Lehours P, Yamaoka Y. Editorial: The Role of Mobile Genetic Elements in Bacterial Evolution and Their Adaptability. Front Microbiol. 2022 Feb 21;13:849667.

7. Arnold BJ, Huang IT, Hanage WP. Horizontal gene transfer and adaptive evolution in bacteria. Nat Rev Microbiol. 2022 Apr;20(4):206–18.

8. Porter SS, Holtappels D, Montoya AP, Koskella B. Causes and consequences of bacterial local adaptation via MGEs in the plant microbiome. New Phytologist. in press;

9. Bottery MJ, Wood AJ, Brockhurst MA. Temporal dynamics of bacteria-plasmid coevolution under antibiotic selection. ISME J. 2019 Feb;13(2):559–62.

10. San Millan A, Heilbron K, MacLean RC. Positive epistasis between co-infecting plasmids promotes plasmid survival in bacterial populations. ISME J. 2014 Mar;8(3):601–12.

11. Wright RCT, Wood AJ, Bottery MJ, Muddiman KJ, Paterson S, Harrison E, et al. A chromosomal mutation is superior to a plasmid-encoded mutation for plasmid fitness cost compensation. PLOS Biology. 2024 Dec 2;22(12):e3002926.

12. Hall JPJ, Brockhurst MA, Harrison E. Sampling the mobile gene pool: innovation via horizontal gene transfer in bacteria. Philosophical Transactions of the Royal Society B: Biological Sciences. 2017 Dec 5;372(1735):20160424.

13. Johnson CN, Sheriff EK, Duerkop BA, Chatterjee A. Let Me Upgrade You: Impact of Mobile Genetic Elements on Enterococcal Adaptation and Evolution. Journal of Bacteriology. 2021 Oct 12;203(21):10.1128/jb.00177-21.

14. Burch CL, Romanchuk A, Kelly M, Wu Y, Jones CD. Empirical Evidence That Complexity Limits Horizontal Gene Transfer. Genome Biology and Evolution. 2023 Jun 1;15(6):evad089.

15. Jain R, Rivera MC, Lake JA. Horizontal gene transfer among genomes: The complexity hypothesis. Proceedings of the National Academy of Sciences. 1999 Mar 30;96(7):3801–6.

16. Novick A, Doolittle WF. Horizontal persistence and the complexity hypothesis. Biol Philos. 2019 Nov 27;35(1):2.

17. Harrison E, Brockhurst MA. Plasmid-mediated horizontal gene transfer is a coevolutionary process. Trends in Microbiology. 2012 Jun;20(6):262–7.

18. Haudiquet M, de Sousa JM, Touchon M, Rocha EPC. Selfish, promiscuous and sometimes useful: how mobile genetic elements drive horizontal gene transfer in microbial populations. Philosophical Transactions of the Royal Society B: Biological Sciences. 2022 Aug 22;377(1861):20210234.

19. Nguyen ANT, Woods LC, Gorrell R, Ramanan S, Kwok T, McDonald MJ. Recombination resolves the cost of horizontal gene transfer in experimental populations of Helicobacter pylori. Proceedings of the National Academy of Sciences. 2022 Mar 22;119(12):e2119010119.

20. Dismukes W, Braga MP, Hembry DH, Heath TA, Landis MJ. Cophylogenetic Methods to Untangle the Evolutionary History of Ecological Interactions. Annual Review of Ecology, Evolution, and Systematics. 2022 Nov 2;53(Volume 53, 2022):275–98.

21. Hayward A, Poulin R, Nakagawa S. A broadscale analysis of host-symbiont cophylogeny reveals the drivers of phylogenetic congruence. Ecology Letters. 2021;24(8):1681–96.

22. Wadgymar SM, DeMarche ML, Josephs EB, Sheth SN, Anderson JT. Local Adaptation: Causal Agents of Selection and Adaptive Trait Divergence. Annual Review of Ecology, Evolution, and Systematics. 2022 Nov 2;53(Volume 53, 2022):87–111.

23. diCenzo GC, Finan TM. The Divided Bacterial Genome: Structure, Function, and Evolution. Microbiology and Molecular Biology Reviews. 2017 Aug 9;81(3):10.1128/mmbr.00019-17.

24. Power JJ, Pinheiro F, Pompei S, Kovacova V, Yüksel M, Rathmann I, et al. Adaptive evolution of hybrid bacteria by horizontal gene transfer. Proceedings of the National Academy of Sciences. 2021 Mar 9;118(10):e2007873118.

25. Bruijning M, Henry LP, Forsberg SKG, Metcalf CJE, Ayroles JF. Natural selection for imprecise vertical transmission in host-microbiota systems. Nat Ecol Evol. 2022 Jan;6(1):77–87.

26. Porter SS. Evolution: Symbiont switching and environmental adaptation. Current Biology. 2021 Sep 13;31(17):R1049–50.

27. Perez-Lamarque B, Morlon H. Distinguishing Cophylogenetic Signal from Phylogenetic Congruence Clarifies the Interplay Between Evolutionary History and Species Interactions. Systematic Biology. 2024 May 1;73(3):613–22.

28. Porter SS, Dupin SE, Denison RF, Kiers ET, Sachs JL. Host-imposed control mechanisms in legume–rhizobia symbiosis. Nat Microbiol. 2024 Aug;9(8):1929–39.

29. Porter SS, Faber-Hammond J, Montoya AP, Friesen ML, Sackos C. Dynamic genomic architecture of mutualistic cooperation in a wild population of Mesorhizobium. The ISME Journal [Internet]. 2018 Sep 14; Available from: 10.1038/s41396-018-0266-y

30. Ramsay JP, Ronson CW. Genetic Regulation of Symbiosis Island Transfer in Mesorhizobium loti. In: Biological Nitrogen Fixation [Internet]. John Wiley & Sons, Ltd; 2015 [cited 2021 Apr 13]. p. 217–24. Available from: http://undefined/doi/abs/10.1002/9781119053095.ch21

31. Sullivan JT, Ronson CW. Evolution of rhizobia by acquisition of a 500-kb symbiosis island that integrates into a phe-tRNA gene. Proc Natl Acad Sci U S A. 1998 Apr 28;95(9):5145– 9.

32. Colombi E, Hill Y, Lines R, Sullivan JT, Kohlmeier MG, Christophersen CT, et al. Population genomics of Australian indigenous Mesorhizobium reveals diverse nonsymbiotic genospecies capable of nitrogen-fixing symbioses following horizontal gene transfer. Microbial Genomics [Internet]. 2023 Jan 5 [cited 2023 Jan 12];9(1). Available from: https://www.microbiologyresearch.org/content/journal/mgen/10.1099/mgen.0.000918

33. Hill YJ, Kohlmeier MG, Agha Amiri A, O’Hara GW, Terpolilli JJ. Evolution of novel Mesorhizobium genospecies that competitively and effectively nodulate Cicer arietinum following inoculation with the Australian commercial inoculant strain M. ciceri CC1192. Plant Soil. 2025 Feb 1;507(1):397–415.

34. Liu S, Jiao J, Tian CF. Adaptive Evolution of Rhizobial Symbiosis beyond Horizontal Gene Transfer: From Genome Innovation to Regulation Reconstruction. Genes. 2023 Feb;14(2):274.

35. Batstone RT, Lindgren H, Allsup CM, Goralka LA, Riley AB, Grillo MA, et al. Genome-Wide Association Studies across Environmental and Genetic Contexts Reveal Complex Genetic Architecture of Symbiotic Extended Phenotypes. mBio. 2022 Oct 26;13(6):e01823–22.

36. Fagorzi C, Bacci G, Huang R, Cangioli L, Checcucci A, Fini M, et al. Nonadditive Transcriptomic Signatures of Genotype-by-Genotype Interactions during the Initiation of Plant-Rhizobium Symbiosis. mSystems. 2021 Jan 12;6(1):10.1128/msystems.00974-20.

37. Riaz MR, Sosa Marquez I, Lindgren H, Levin G, Doyle R, Romero MC, et al. Mobile gene clusters and coexpressed plant–rhizobium pathways drive partner quality variation in symbiosis. Proceedings of the National Academy of Sciences. 2025 Aug 5;122(31):e2411831122.

38. Boivin S, Ait Lahmidi N, Sherlock D, Bonhomme M, Dijon D, Heulin-Gotty K, et al. Host-specific competitiveness to form nodules in Rhizobium leguminosarum symbiovar viciae. New Phytol. 2020 Apr;226(2):555–68.

39. Perry BJ, Sullivan JT, Colombi E, Murphy RJT, Ramsay JP, Ronson CW. Symbiosis islands of Loteae-nodulating Mesorhizobium comprise three radiating lineages with concordant nod gene complements and nodulation host-range groupings. Microb Genom. 2020 Aug 26;6(9):mgen000426.

40. Roche P, Maillet F, Plazanet C, Debellé F, Ferro M, Truchet G, et al. The common nodABC genes of Rhizobium meliloti are host-range determinants. Proc Natl Acad Sci U S A. 1996 Dec 24;93(26):15305–10.

41. Fagorzi C, Checcucci A, DiCenzo GC, Debiec-Andrzejewska K, Dziewit L, Pini F, et al. Harnessing Rhizobia to Improve Heavy-Metal Phytoremediation by Legumes. Genes. 2018 Nov;9(11):542.

42. Kehlet-Delgado H, Montoya AP, Jensen KT, Wendlandt CE, Dexheimer C, Roberts M, et al. The evolutionary genomics of adaptation to stress in wild rhizobium bacteria. Proceedings of the National Academy of Sciences. 2024 Mar 26;121(13):e2311127121.

43. Porter SS, Chang PL, Conow CA, Dunham JP, Friesen ML. Association mapping reveals novel serpentine adaptation gene clusters in a population of symbiotic Mesorhizobium. ISME J. 2017 Jan;11(1):248–62.

44. Jensen KT, Griffitts JS. Regulation of a nickel tolerance operon conserved in Mesorhizobium strains from serpentine soils [Internet]. bioRxiv; 2025 [cited 2025 Oct 16]. p. 2025.07.11.664438. Available from: https://www.biorxiv.org/content/10.1101/2025.07.11.664438v1

45. Audrey B, Cellier N, White F, Jacques PÉ, Burrus V. A systematic approach to classify and characterize genomic islands driven by conjugative mobility using protein signatures. Nucleic Acids Research. 2023 Sep 8;51(16):8402–12.

46. Liu M, Li X, Xie Y, Bi D, Sun J, Li J, et al. ICEberg 2.0: an updated database of bacterial integrative and conjugative elements. Nucleic Acids Research. 2019 Jan 8;47(D1):D660–5.

47. Cury J, Touchon M, Rocha EPC. Integrative and conjugative elements and their hosts: composition, distribution and organization. Nucleic Acids Res. 2017 Sep 6;45(15):8943–56.

48. Rocha EPC, Danchin A. Base composition bias might result from competition for metabolic resources. Trends in Genetics. 2002 Jun 1;18(6):291–4.

49. Brockhurst MA, Cameron DD, Beckerman AP. Fitness trade-offs and the origins of endosymbiosis. PLOS Biology. 2024 Apr 12;22(4):e3002580.

50. Blasco-Costa I, Hayward A, Poulin R, Balbuena JA. Next-generation cophylogeny: unravelling eco-evolutionary processes. Trends in Ecology & Evolution. 2021 Oct 1;36(10):907–18.

51. Alderliesten JB, Duxbury SJN, Zwart MP, de Visser JAGM, Stegeman A, Fischer EAJ. Effect of donor-recipient relatedness on the plasmid conjugation frequency: a meta-analysis. BMC Microbiol. 2020 May 26;20(1):135.

52. Porse A, Schou TS, Munck C, Ellabaan MMH, Sommer MOA. Biochemical mechanisms determine the functional compatibility of heterologous genes. Nat Commun. 2018 Feb 6;9(1):522.

53. Ares-Arroyo M, Coluzzi C, Sousa JAM de, Rocha EPC. Hijackers, hitchhikers, or co-drivers? The mysteries of mobilizable genetic elements. PLOS Biology. 2024 Aug 29;22(8):e3002796.

54. Burghardt LT, diCenzo GC. The evolutionary ecology of rhizobia: multiple facets of competition before, during, and after symbiosis with legumes. Current Opinion in Microbiology. 2023 Apr 1;72:102281.

55. Wheatley RM, Ford BL, Li L, Aroney STN, Knights HE, Ledermann R, et al. Lifestyle adaptations of Rhizobium from rhizosphere to symbiosis. Proceedings of the National Academy of Sciences of the United States of America. 2020 Sep 8;117(38):23823.

56. Baiotto T, Guzman LM. Maintaining Local Adaptation Is Key for Evolutionary Rescue and Long-Term Persistence of Populations Experiencing Habitat Loss and a Changing Environment. Evolutionary Applications. 2025;18(3):e70081.

57. Ferenci T. Trade-off Mechanisms Shaping the Diversity of Bacteria. Trends Microbiol. 2016 Mar;24(3):209–23.

58. Bottery MJ, Wood AJ, Brockhurst MA. Adaptive modulation of antibiotic resistance through intragenomic coevolution. Nat Ecol Evol. 2017 Sep;1(9):1364–9.

59. Santichaivekin S, Yang Q, Liu J, Mawhorter R, Jiang J, Wesley T, et al. eMPRess: a systematic cophylogeny reconciliation tool. Bioinformatics. 2021 Aug 25;37(16):2481–2.

60. Pérez Carrascal OM, VanInsberghe D, Juárez S, Polz MF, Vinuesa P, González V. Population genomics of the symbiotic plasmids of sympatric nitrogen-fixing Rhizobium species associated with Phaseolus vulgaris. Environmental Microbiology. 2016;18(8):2660– 76.

61. Riley AB, Grillo MA, Epstein B, Tiffin P, Heath KD. Discordant population structure among rhizobium divided genomes and their legume hosts. Molecular Ecology. 2023;32(10):2646–59.

62. Wang X, Liu D, Luo Y, Zhao L, Liu Z, Chou M, et al. Comparative analysis of rhizobial chromosomes and plasmids to estimate their evolutionary relationships. Plasmid. 2018 Mar 1;96–97:13–24.

63. Davies-Barnard T, Friedlingstein P. The Global Distribution of Biological Nitrogen Fixation in Terrestrial Natural Ecosystems. Global Biogeochemical Cycles. 2020;34(3):e2019GB006387.

64. Greenlon A, Chang PL, Damtew ZM, Muleta A, Carrasquilla-Garcia N, Kim D, et al. Global-level population genomics reveals differential effects of geography and phylogeny on horizontal gene transfer in soil bacteria. PNAS. 2019 Jul 23;116(30):15200–9.

65. Karasev ES, Hosid SL, Aksenova TS, Onishchuk OP, Kurchak ON, Dzyubenko NI, et al. Impacts of Natural Selection on Evolution of Core and Symbiotically Specialized (sym) Genes in the Polytypic Species Neorhizobium galegae. International Journal of Molecular Sciences. 2023 Jan;24(23):16696.

66. Andrews M, De Meyer S, James EK, Stępkowski T, Hodge S, Simon MF, et al. Horizontal Transfer of Symbiosis Genes within and Between Rhizobial Genera: Occurrence and Importance. Genes (Basel). 2018 Jun 27;9(7):321.

67. Gunununu RP, Mohammed M, Jaiswal SK, Dakora FD. Phylogeny and symbiotic effectiveness of indigenous rhizobial microsymbionts of common bean (Phaseolus vulgaris L.) in Malkerns, Eswatini. Sci Rep. 2023 Oct 9;13(1):17029.

68. Weisberg AJ, Pearce E, Kramer CG, Chang JH, Clarke CR. Diverse mobile genetic elements shaped the evolution of Streptomyces virulence. Microbial Genomics. 2023;9(11):001127.

69. Remigi P, Zhu J, Young JPW, Masson-Boivin C. Symbiosis within Symbiosis: Evolving Nitrogen-Fixing Legume Symbionts. Trends in Microbiology. 2016 Jan;24(1):63–75.

70. Wardell GE, Hynes MF, Young PJ, Harrison E. Why are rhizobial symbiosis genes mobile? Philosophical Transactions of the Royal Society B: Biological Sciences. 2022 Jan 17;377(1842):20200471.

71. Andersson DI, Hughes D. Antibiotic resistance and its cost: is it possible to reverse resistance? Nat Rev Microbiol. 2010 Apr;8(4):260–71.

72. Frederickson ME. Rethinking Mutualism Stability: Cheaters and the Evolution of Sanctions. The Quarterly Review of Biology. 2013 Dec;88(4):269–95.

73. Friesen ML. Widespread fitness alignment in the legume–rhizobium symbiosis. New Phytologist. 2012;194(4):1096–111.

74. Sachs JL, Russell JE, Hollowell AC. Evolutionary Instability of Symbiotic Function in Bradyrhizobium japonicum. PLOS ONE. 2011 Nov 2;6(11):e26370.

75. Geddes BA, Kearsley J, Morton R, diCenzo GC, Finan TM. Chapter Eight - The genomes of rhizobia. In: Frendo P, Frugier F, Masson-Boivin C, editors. Advances in Botanical Research [Internet]. Academic Press; 2020 [cited 2024 Oct 20]. p. 213–49. (Regulation of Nitrogen-Fixing Symbioses in Legumes; vol. 94). Available from: https://www.sciencedirect.com/science/article/pii/S0065229619300916

76. Milunovic B, diCenzo GC, Morton RA, Finan TM. Cell Growth Inhibition upon Deletion of Four Toxin-Antitoxin Loci from the Megaplasmids of Sinorhizobium meliloti. Journal of Bacteriology. 2014 Jan 23;196(4):811–24.

77. diCenzo GC, MacLean AM, Milunovic B, Golding GB, Finan TM. Examination of Prokaryotic Multipartite Genome Evolution through Experimental Genome Reduction. PLOS Genetics. 2014 Oct 23;10(10):e1004742.

78. Weisberg AJ, Rahman A, Backus D, Tyavanagimatt P, Chang JH, Sachs JL. Pangenome Evolution Reconciles Robustness and Instability of Rhizobial Symbiosis. mBio. 2022 Apr 13;13(3):e00074–22.

79. Touchon M, Bobay LM, Rocha EP. The chromosomal accommodation and domestication of mobile genetic elements. Current Opinion in Microbiology. 2014 Dec 1;22:22–9.

80. Hao C, Dewar AE, West SA, Ghoul M. Gene transferability and sociality do not correlate with gene connectivity. Proceedings of the Royal Society B: Biological Sciences. 2022 Nov 30;289(1987):20221819.

81. Wang Y, Batra A, Schulenburg H, Dagan T. Gene sharing among plasmids and chromosomes reveals barriers for antibiotic resistance gene transfer. Philosophical Transactions of the Royal Society B: Biological Sciences. 2021 Nov 29;377(1842):20200467.

82. Bañuelos-Vazquez LA, Castellani LG, Luchetti A, Romero D, Tejerizo GAT, Brom S. Role of plant compounds in the modulation of the conjugative transfer of pRet42a. PLOS ONE. 2020 Aug 26;15(8):e0238218.

83. Ling J, Wang H, Wu P, Li T, Tang Y, Naseer N, et al. Plant nodulation inducers enhance horizontal gene transfer of Azorhizobium caulinodans symbiosis island. Proc Natl Acad Sci U S A. 2016 Nov 29;113(48):13875–80.

84. Somasegaran P, Hoben HJ. Handbook for Rhizobia: Methods in Legume-Rhizobium Technology [Internet]. New York: Springer-Verlag; 1994 [cited 2021 Apr 14]. Available from: https://www.springer.com/us/book/9781461383772

85. Wendlandt CE, Helliwell E, Roberts M, Nguyen KT, Friesen ML, von Wettberg E, et al. Decreased coevolutionary potential and increased symbiont fecundity during the biological invasion of a legume-rhizobium mutualism. Evolution. 2021;75(3):731–47.

86. Darling AE, Mau B, Perna NT. progressiveMauve: Multiple Genome Alignment with Gene Gain, Loss and Rearrangement. PLOS ONE. 2010 Jun 25;5(6):e11147.

87. Grant JR, Enns E, Marinier E, Mandal A, Herman EK, Chen C yu, et al. Proksee: in-depth characterization and visualization of bacterial genomes. Nucleic Acids Research. 2023 Jul 5;51(W1):W484–92.

88. Bertelli C, Laird MR, Williams KP, Lau BY, Hoad G, Winsor GL, et al. IslandViewer 4: expanded prediction of genomic islands for larger-scale datasets. Nucleic Acids Res. 2017 Jul 3;45(Web Server issue):W30–5.

89. de Jong A, Kuipers OP, Kok J. FUNAGE-Pro: comprehensive web server for gene set enrichment analysis of prokaryotes. Nucleic Acids Res. 2022 May 31;50(W1):W330–6.

90. Néron B, Denise R, Coluzzi C, Touchon M, Rocha EPC, Abby SS. MacSyFinder v2: Improved modelling and search engine to identify molecular systems in genomes. Peer Community Journal [Internet]. 2023 [cited 2025 Mar 21];3. Available from: https://peercommunityjournal.org/articles/10.24072/pcjournal.250/

91. Tesson F, Hervé A, Mordret E, Touchon M, d’Humières C, Cury J, et al. Systematic and quantitative view of the antiviral arsenal of prokaryotes. Nat Commun. 2022 May 10;13(1):2561.

92. Alcock BP, Huynh W, Chalil R, Smith KW, Raphenya AR, Wlodarski MA, et al. CARD 2023: expanded curation, support for machine learning, and resistome prediction at the Comprehensive Antibiotic Resistance Database. Nucleic Acids Research. 2023 Jan 6;51(D1):D690–9.

93. Arndt D, Grant JR, Marcu A, Sajed T, Pon A, Liang Y, et al. PHASTER: a better, faster version of the PHAST phage search tool. Nucleic Acids Research. 2016 Jul 8;44(W1):W16–21.

94. Camargo AP, Roux S, Schulz F, Babinski M, Xu Y, Hu B, et al. Identification of mobile genetic elements with geNomad. Nat Biotechnol. 2024 Aug;42(8):1303–12.

95. Bates D, Mächler M, Bolker B, Walker S. Fitting Linear Mixed-Effects Models Using lme4. Journal of Statistical Software. 2015 Oct 7;67:1–48.

96. Lenth RV, Banfai B, Bolker B, Buerkner P, Giné-Vázquez I, Herve M, et al. emmeans: Estimated Marginal Means, aka Least-Squares Means [Internet]. 2025 [cited 2025 Jun 17]. Available from: https://cran.r-project.org/web/packages/emmeans/index.html

97. Searle SR, Speed FM, Milliken GA. Population Marginal Means in the Linear Model: An Alternative to Least Squares Means. The American Statistician. 1980 Nov 1;34(4):216–21.

98. Hartig F. DHARMa: Residual Diagnostics for Hierarchical (Multi-Level / Mixed) Regression Models [Internet]. 2016 [cited 2024 Oct 23]. p. 0.4.7. Available from: https://CRAN.R-project.org/package=DHARMa

99. R Core Team. R: A language and environment for statistical computing. R Foundation for Statistical Computing, Vienna, Austria. 2022; Available from: https://www.R-project.org/.

100. Stamatakis A. RAxML version 8: a tool for phylogenetic analysis and post-analysis of large phylogenies. Bioinformatics. 2014 May 1;30(9):1312–3.

101. Paradis E, Schliep K. ape 5.0: an environment for modern phylogenetics and evolutionary analyses in R. Bioinformatics. 2019 Feb 1;35(3):526–8.

102. Yu G, Smith DK, Zhu H, Guan Y, Lam TTY. ggtree: an r package for visualization and annotation of phylogenetic trees with their covariates and other associated data. Methods in Ecology and Evolution. 2017;8(1):28–36.

103. Jain C, Rodriguez-R LM, Phillippy AM, Konstantinidis KT, Aluru S. High throughput ANI analysis of 90K prokaryotic genomes reveals clear species boundaries. Nat Commun. 2018 Nov 30;9(1):5114.

104. Balbuena JA, Míguez-Lozano R, Blasco-Costa I. PACo: A Novel Procrustes Application to Cophylogenetic Analysis. PLOS ONE. 2013 Apr 8;8(4):e61048.

105. Hutchinson MC, Cagua EF, Balbuena JA, Stouffer DB, Poisot T. paco: implementing Procrustean Approach to Cophylogeny in R. Methods in Ecology and Evolution. 2017;8(8):932–40.

106. Horne T, Orr VT, Hall JP. How do interactions between mobile genetic elements affect horizontal gene transfer? Current Opinion in Microbiology. 2023 Jun 1;73:102282.

107. Oksanen J, Simpson GL, Blanchet FG, Kindt R, Legendre P, Minchin PR, et al. vegan: Community Ecology Package [Internet]. 2022 [cited 2023 Nov 29]. Available from: https://cran.r-project.org/web/packages/vegan/index.html

108. Rice P, Longden I, Bleasby A. EMBOSS: The European Molecular Biology Open Software Suite. Trends in Genetics. 2000 Jun 1;16(6):276–7.

109. Kassambara A. rstatix: Pipe-Friendly Framework for Basic Statistical Tests. R package version 0.7.2. 2023; Available from: https://rpkgs.datanovia.com/rstatix/.

110. Revell LJ. phytools 2.0: an updated R ecosystem for phylogenetic comparative methods (and other things). PeerJ. 2024 Jan 5;12:e16505.

