## Supplemental Information for "Mobile genetic elements shape the evolution of wild bacterial pangenomes in the face of biotic and abiotic selective pressures in the soil"

This file includes supporting text, figures S1 to S7, and tables S12 and S13.

### I. Integrative plasmid construction to track the *Mesorhizobium* nickel resistance island (NRI)

A plasmid was designed to track the NRI during conjugative transfer and contains selectable markers and a region of homology with the NRI to enable the plasmid to integrate site-specifically into the NRI via homologous recombination (Figure S1). We targeted a 500 bp region at the end of the *ODH* gene (including the stop codon and intergenic region after the gene) in the NRI for the region of homology. The NRI tracking plasmid (pKJ115) was first introduced via bacterial transformation into specialized *E. coli* strain MFDpir which is auxotrophic for DAP (diaminopimelic acid- a lysine precursor). MFDpir isolates with the reporter plasmid (pKJ115) were grown at 37°C overnight on (LB) agar (1) supplemented with 50 µg/mL DAP and 30 µg/mL Kanamycin. *Mesorhizobium* strain C089B was grown at 28°C for 5 days on Tryptone Yeast (TY) agar (2) Approximately equal amounts of MFDpir and *Mesorhizobium* cells were re-suspended in LB broth or TY broth, respectively, and 12 µL of each donor and recipient were mixed in separate 1.5 mL microcentrifuge tubes. 20µL of each bi-parental mating mixture was spot plated on TY agar with 50 µg/mL DAP and incubated at 28°C for 6 hours prior to selecting for *Mesorhizobium* NRI donor transconjugants on TY agar with 100 µg/mL neomycin. Then the *Mesorhizobium* NRI donor transconjugant, P15089B, was cryopreserved after testing on X-Gluc media, where transconjugants with the tracking plasmid turn blue due to the presence of the *GUS* gene.

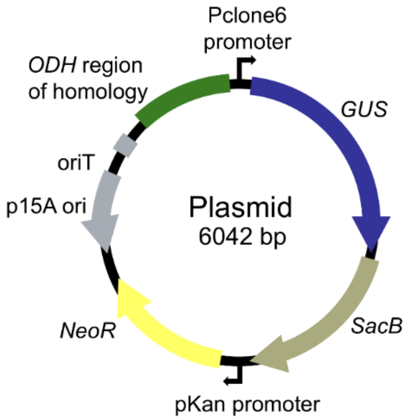


**Figure S1. Genetic map of the nickel resistance island (NRI) tracking plasmid that inserts into the NRI of a *Mesorhizobium* donor strain.** The *Mesorhizobium* NRI in strain C089B is marked with a plasmid (pKJ115) containing selectable marker genes for neomycin resistance (NeoR), *GUS* (beta-glucuronidase), *SacB* (not used in this experiment), and a region of homology with the *ODH* gene to enable the plasmid to integrate site specifically into the intergenic region after the *ODH* gene via homologous recombination. The map indicates locations of the promoters, *Pclone6* and *pKan*, reporter gene (*GUS*), the kanamycin/neomycin resistance gene (*NeoR/KanR*), the cis-acting mobilization determinant (*oriT*), and the *E. coli* replication origin, *p15A ori*.

**II: NRI transfer and function**

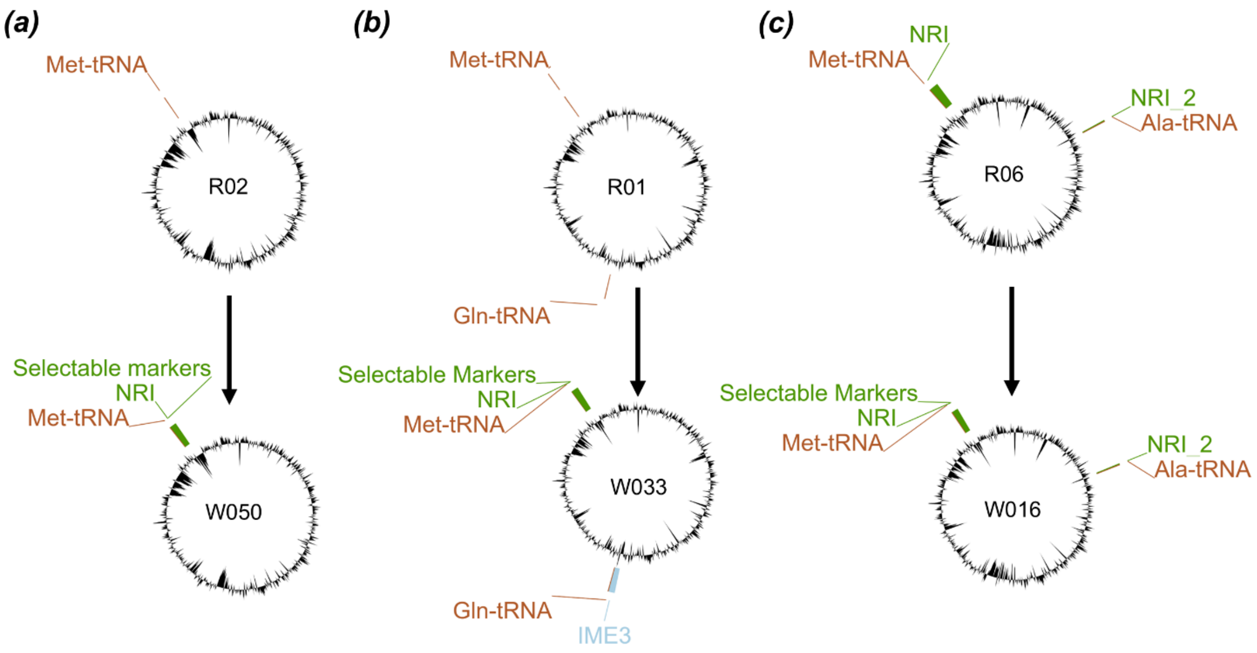


**Figure S2. Genomic maps of recipients and transconjugants from the NRI transmission and growth assays.** Genomic maps made with proksee (3) show the recipient (top) and respective transconjugant (bottom). (a) Recipient R02 has a putative genomic island at the Met-tRNA gene insertion site that moved downstream of the NRI upon its insertion, (b) Recipient R01 gained the NRI at the Met-tRNA gene and another putative integrative and mobilizable element (IME3) at a Gln-tRNA gene from the donor. (c) Recipient R06 has two NRI regions, one inserted at the Met-tRNA gene and another at an Ala-tRNA gene. The original NRI at the Met-tRNA was replaced by the donor NRI. See Table S1 for additional details.

***Examination of MGEs in the NRI donor***

In the NRI donor, C089B, we explored putative MGEs and their gene content. Defense-finder (4) identified a putative antidefense system in ICE2, *ardC*, an anti-restriction modification system that inhibits recipients with type I restriction modification systems from degrading a novel, foreign MGE (5) (Table S3-S5). It also identified the bNACHT Hypnos system on IME3, a putative anti-phage defense system observed in other *Mesorhizobium* (6), and homologs of *GajB* in IME1 and IME4, which is part of a putative Gabija anti-phage system (7) (Table S3-S5). CARD RGI (8) returned one strict hit in ICE1 for a putative antibiotic efflux homolog (Table S6). FUNGAGE-Pro (9) functional enrichment analysis revealed that both ICE2 and IME1 are enriched in gene ontology (GO) terms related to metabolite binding and transport into or out of the cell, such as ATP-binding cassette (ABC) transporter complexes (GO:0043190) (Table S7-

S8). IME1 is enriched in photoreceptor activity (GO:0009881) and contains a putative blue-light activated protein and blue-light histidine kinase (LOV-HK) (Table S8). These genes may function similarly to R-LOV-HK, which some *Rhizobium leguminosarum* carry on their chromid. This locus regulates root attachment during nodulation and the formation of flagella or exopolysaccharides in response to light (10), and may enhance competitiveness for nodulation in *Bradyrhizobium* (11).

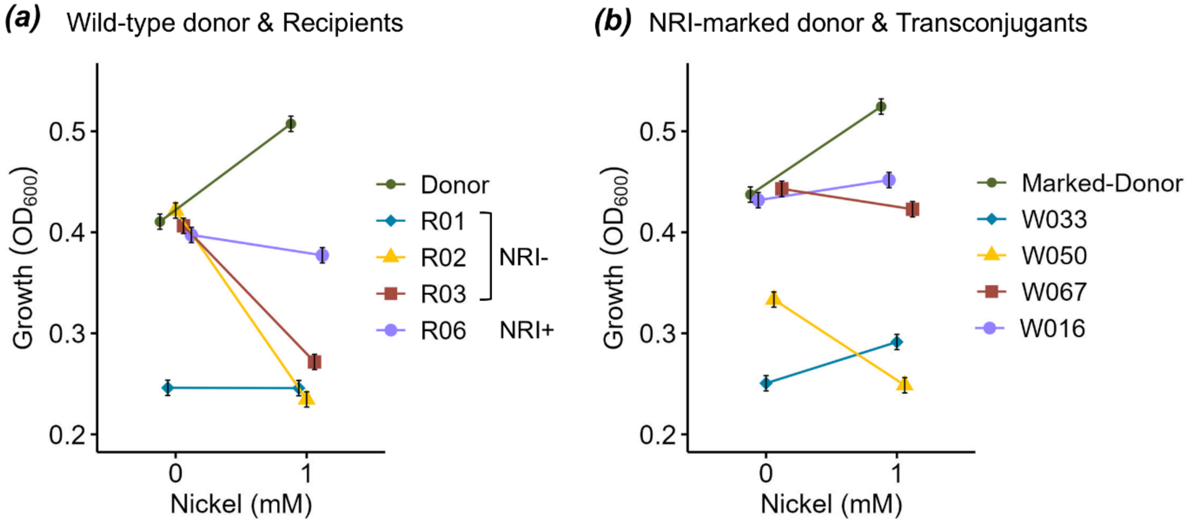


**Figure S3. Growth reaction norms in the presence or absence of nickel.** The estimated marginal mean growth ($OD_{600}$ at 72 hours) +/- standard error of (a) original strains (wild-type donor and recipients) and (b) the NRI-marked donor and NRI gain transconjugants. The same shape and color points are used for pairs of strains to compare (i.e., the wild-type donor and transgenic donor bearing a marker plasmid inserted in the NRI or the recipients and respective transconjugants). Estimated marginal means (12,13) are derived from model of growth predicted by the interaction of strain identity and nickel level ($\chi 2 = 233$, $P < 0.001$).

### III. Genomic compartment characteristics and ecological variables

As expected, the type of soil (serpentine or non-serpentine) a strain was collected from predicted MIC and soil nickel in the NRI model (Wilks' lambda = 0.53, $F_{2, 253} = 113.5$, $P < 0.001$; Figure S4c), SI model (Wilks' lambda = 0.33, $F_{2, 284} = 288.3$, $P < 0.001$; Figure 2h), and chromosome model (Wilks' lambda = 0.33, $F_{2, 281} = 285.7$, $P < 0.001$; Figure S6). This allowed us to better distinguish MGE/chromosome co-inheritance patterns in particular soil types from phylotype or clade associations with nickel MIC and soil nickel level.

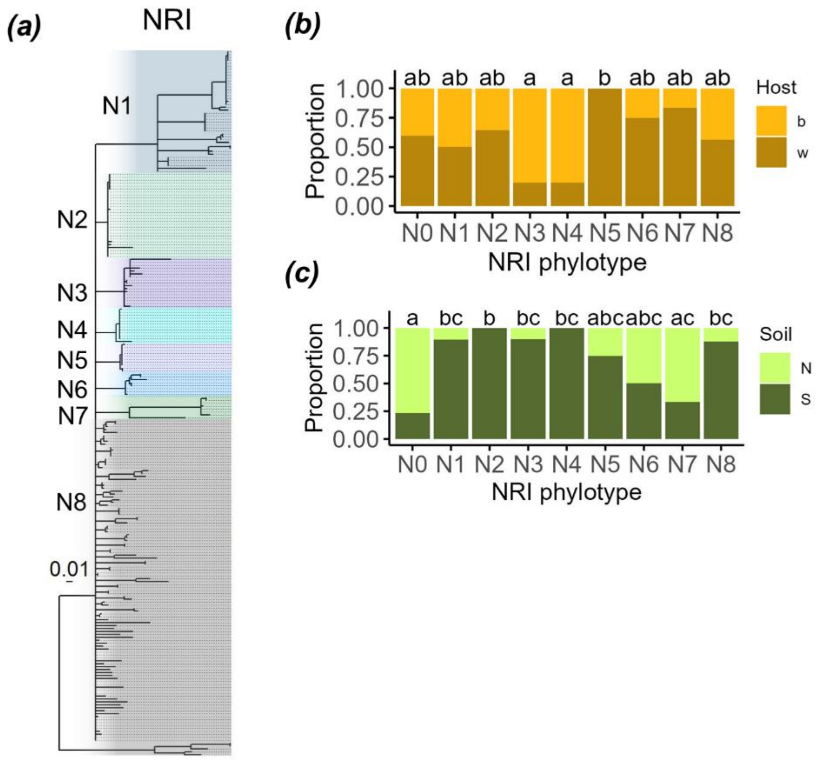


**Figure S4. NRI phylogeny and phylotype designations.** (a) NRI phylogeny based on amino acid sequences of *nreA* and *nreX* genes built with RAxML. Branches with bootstrap support values of less than 70 are collapsed to polytomies. Scale bar indicates substitutions per site. The proportion of (b) host type (*Acmispon brachycarpus* "b", *A.* wrangelianus "w"), and (c) soil type (non-serpentine "N", serpentine "S") bearing each NRI phylotype. The association found between host plant species and NRI phylotype appears driven by the observance that NRI phylotype N5 (n = 8) only associate with strains from *A. brachycarpus*, while phylotypes N3 (n = 10) and N4 (n = 10) associate with *A. wrangelianus* in 80% of observations.

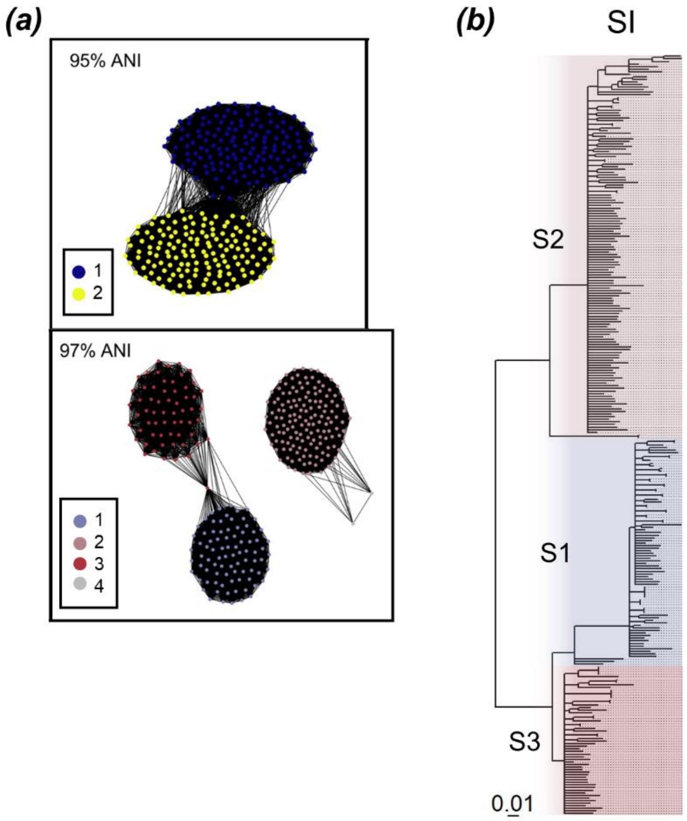


**Figure S5. The core symbiosis island (SI) phylotypes were defined using more distinct clusters at 97% average nucleotide identity (ANI) compared to 95% ANI**. (a) Weighted undirected graphs were constructed using pairwise average nucleotide identity (ANI) values calculated among 295 *Mesorhizobium* genomes that contain the SI. ANI was calculated from 21 single-copy core genes in the SI. Each node represents a genome and the weighted edges connecting nodes represent a pairwise distance ANI value of 95% (top) or 97% (bottom). Nodes are colored by the medoid genome of the cluster formed at the 95% or 97% ANI value. Cluster 4 only contains two strains and was incorporated into cluster 2. Graphs were constructed using the R (v 4.2.2) (14) package bactaxR (v 0.2.3) (15). (b) SI phylogeny based on amino acid sequences of 21 single-copy core genes built with RAxML. Branches with bootstrap support values of less than 70 are collapsed to polytomies. Scale bar indicates substitutions per site.

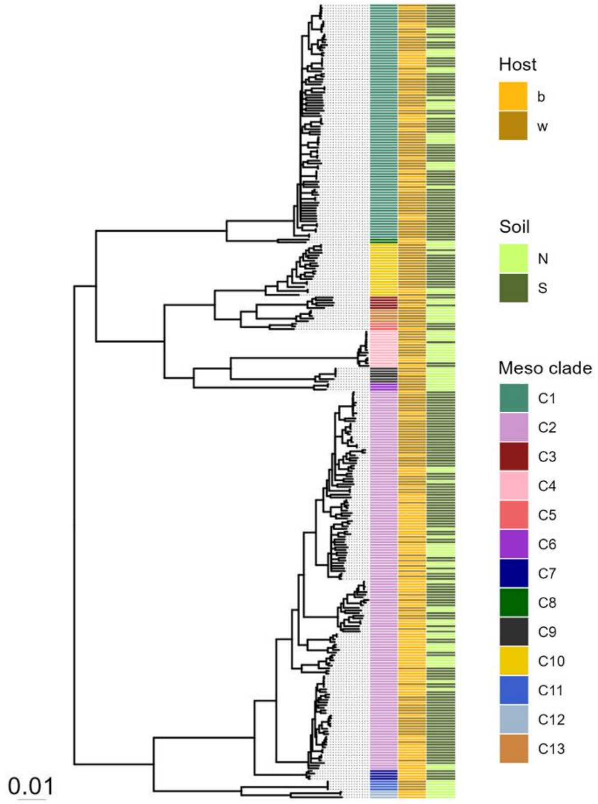


**Figure S6. The core chromosome is non-randomly associated with host plant species.** *Mesorhizobium* chromosome phylogeny based on amino acid sequences of 1,542 single copy core genes built with RAxML (16). The *Mesorhizobium* clade (C1-C13), host plant (*Acmispon brachycarpus* "b", *A. wrangelianus* "w"), and soil type (non-serpentine "N", serpentine "S") for each strain are displayed as colored tiles. Branches with bootstrap support values of less than 70 are collapsed to polytomies. Scale bar indicates substitutions per site.

**Table S12. Mantel correlation tests for the chromosome and MGEs.** Mantel tests compare geographic distance and phylogenetic distance of the chromosome or MGEs, as well as between the core chromosome phylogenetic distance and MGE phylogenetic distance, and between MGEs.

| | Geographic distance | | Core chromosome phylogenetic distance | | Core SI phylogenetic distance | |
|---|---|---|---|---|---|---|
| Component | Mantel *r* | *p* | Mantel *r* | *p* | Mantel *r* | *p* |
| Chromosome | 0.005 | 0.39 | . | . | . | . |
| SI | 0.007 | 0.25 | 0.92 | 0.001*** | . | . |
| NRI | 0.022 | 0.29 | 0.28 | 0.001*** | 0.24 | 0.001*** |

**Table S13. Estimates of nucleotide diversity and GC content of each genomic compartment.** Calculations were performed using the core genes of each genomic compartment and estimates reflect the average among strains. For the NRI, we excluded strains with more than one copy of *nreAX*, but estimates were similar regardless.

| Genomic compartment | Nucleotide diversity ($\pi$) | GC% |
|---|---|---|
| Chromosome | 0.048 | 59.4 |
| SI | 0.021 | 60.5 |
| NRI | 0.065 | 62 |

**IV. NRI ancestral state reconstruction**

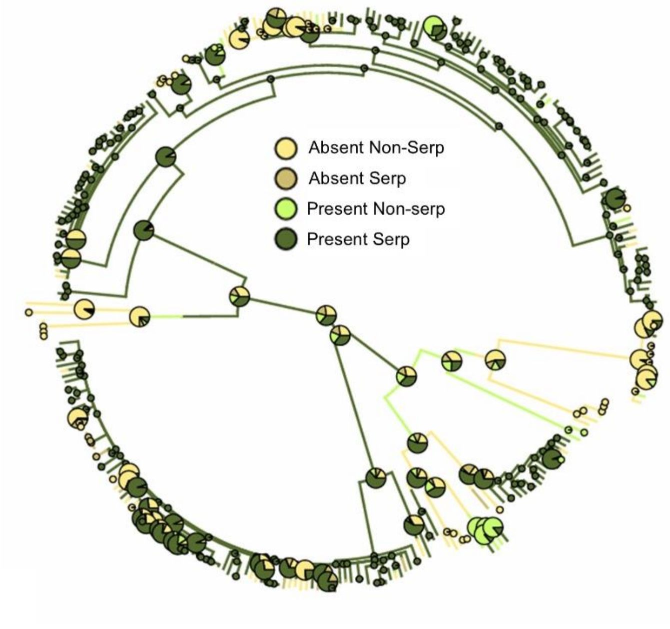


**Figure S7. Posterior probabilities at each ancestral node of the *Mesorhizobium* phylogeny for the ancestral state reconstruction of NRI character states based on the presence or absence of NRI on serpentine or non-serpentine soils.** A randomly selected map from the 1000 stochastic maps with posterior probabilities overlaid at nodes from R package phytools (v 2.1-1) (17). The posterior probabilities from stochastic character mapping that each node is in each state are shown at internal nodes and the observed discrete character states are shown at the tips of the tree. Internal nodes in which no state has 95% support are larger than less ambiguous nodes.

**Table S14. Transition frequencies and Bayesian 95% high probability density (HPD) intervals across the 1000 stochastic maps of the ancestral state reconstruction of NRI character states based on the presence or absence of NRI on serpentine or non-serpentine soils.** The HPD indicates the 95% high probability density interval for transitions between each NRI character state.

| Transition type | Mean transition frequencies | HPD interval |
|---|---|---|
| Present Serp → Absent Non-serp | 27.2 | [20, 33] |
| Present Serp → Absent Serp | 15.1 | [11, 19] |
| Absent Non-serp → Present Serp | 12.9 | [7, 18] |
| Present Serp → Present Non-serp | 9.9 | [6, 13] |
| Absent Non-serp → Present Non-serp | 6.6 | [4, 9] |
| Absent Non-serp → Absent Serp | 4.9 | [2, 8] |
| Present Non-serp → Present Serp | 4.7 | [2, 8] |
| Absent Serp → Absent Non-serp | 2.9 | [0, 6] |
| Present Non-serp → Absent Non-serp | 2.7 | [0, 6] |
| Absent Serp → Present Serp | 2.6 | [0, 6] |
| Present Non-serp → Absent Serp | 1.14 | [0, 3] |
| Absent Serp → Present Non-serp | 1.12 | [0, 3] |

### SUPPLEMENTAL REFERENCES


1. Bertani G. STUDIES ON LYSOGENESIS I.: The Mode of Phage Liberation by Lysogenic Escherichia coli. Journal of Bacteriology. 1951 Sep 1;62(3):293–300.

2. Somasegaran P, Hoben HJ. Handbook for Rhizobia: Methods in Legume-Rhizobium Technology [Internet]. New York: Springer-Verlag; 1994 [cited 2021 Apr 14]. Available from: https://www.springer.com/us/book/9781461383772

3. Grant JR, Enns E, Marinier E, Mandal A, Herman EK, Chen C yu, et al. Proksee: in-depth characterization and visualization of bacterial genomes. Nucleic Acids Research. 2023 Jul 5;51(W1):W484–92.

4. Tesson F, Planel R, Egorov AA, Georjon H, Vaysset H, Brancotte B, et al. A Comprehensive Resource for Exploring Antiphage Defense: DefenseFinder Webservice,Wiki and Databases. Peer Community Journal [Internet]. 2024 [cited 2025 Mar 21];4. Available from: https://peercommunityjournal.org/articles/10.24072/pcjournal.470/

5. González-Montes L, del Campo I, Garcillán-Barcia MP, de la Cruz F, Moncalián G. ArdC, a ssDNA-binding protein with a metalloprotease domain, overpasses the recipient hsdRMS restriction system broadening conjugation host range. PLoS Genet. 2020 Apr 29;16(4):e1008750.

6. Berg DF van den, Costa AR, Esser JQ, Stanciu I, Geissler JQ, Zoumaro-Djayoon AD, et al. Bacterial homologs of innate eukaryotic antiviral defenses with anti-phage activity highlight shared evolutionary roots of viral defenses. Cell Host & Microbe. 2024 Aug 14;32(8):1427-1443.e8.

7. Huo Y, Kong L, Zhang Y, Xiao M, Du K, Xu S, et al. Structural and biochemical insights into the mechanism of the Gabija bacterial immunity system. Nat Commun. 2024 Jan 29;15:836.

8. Alcock BP, Huynh W, Chalil R, Smith KW, Raphenya AR, Wlodarski MA, et al. CARD 2023: expanded curation, support for machine learning, and resistome prediction at the Comprehensive Antibiotic Resistance Database. Nucleic Acids Research. 2023 Jan 6;51(D1):D690–9.

9. de Jong A, Kuipers OP, Kok J. FUNAGE-Pro: comprehensive web server for gene set enrichment analysis of prokaryotes. Nucleic Acids Res. 2022 May 31;50(W1):W330–6.

10. Bonomi HR, Posadas DM, Paris G, Carrica M del C, Frederickson M, Pietrasanta LI, et al. Light regulates attachment, exopolysaccharide production, and nodulation in Rhizobium leguminosarum through a LOV-histidine kinase photoreceptor. Proceedings of the National Academy of Sciences. 2012 Jul 24;109(30):12135–40.

11. Rahman A, Manci M, Nadon C, Perez IA, Farsamin WF, Lampe MT, et al. Competitive interference among rhizobia reduces benefits to hosts. Current Biology. 2023 Jul 24;33(14):2988-3001.e4.

12. Lenth RV, Banfai B, Bolker B, Buerkner P, Giné-Vázquez I, Herve M, et al. emmeans: Estimated Marginal Means, aka Least-Squares Means [Internet]. 2025 [cited 2025 Jun 17]. Available from: https://cran.r-project.org/web/packages/emmeans/index.html

13. Searle SR, Speed FM, Milliken GA. Population Marginal Means in the Linear Model: An Alternative to Least Squares Means. The American Statistician. 1980 Nov 1;34(4):216–21.

14. R Core Team. R: A language and environment for statistical computing. R Foundation for Statistical Computing, Vienna, Austria. 2022; Available from: https://www.R-project.org/.

15. Carroll LM, Wiedmann M, Kovac J. Proposal of a Taxonomic Nomenclature for the Bacillus cereus Group Which Reconciles Genomic Definitions of Bacterial Species with Clinical and Industrial Phenotypes. mBio. 2020 Feb 25;11(1):10.1128/mbio.00034-20.

16. Kehlet-Delgado H, Montoya AP, Jensen KT, Wendlandt CE, Dexheimer C, Roberts M, et al. The evolutionary genomics of adaptation to stress in wild rhizobium bacteria. Proceedings of the National Academy of Sciences. 2024 Mar 26;121(13):e2311127121.

17. Revell LJ. phytools 2.0: an updated R ecosystem for phylogenetic comparative methods (and other things). PeerJ. 2024 Jan 5;12:e16505.
